# A hierarchical coordinate system for sequence memory in human entorhinal cortex

**DOI:** 10.1101/2024.10.30.620612

**Authors:** Anna Shpektor, Jacob J. W. Bakermans, Alon B. Baram, Johannes Sarnthein, Debora Ledergerber, Lukas Imbach, Alina Kiseleva, Emma Müller-Seydlitz, Helen C. Barron, Timothy E. J. Behrens

**Affiliations:** Wellcome Centre for Integrative Neuroimaging, University of Oxford, Oxford, UK; Sainsbury Wellcome Centre for Neural Circuits and Behaviour, University College London, London, UK; Klinik für Neurochirurgie, Universitätsspital Zürich, Universität Zürich, Zurich, Switzerland; Zurich Neuroscience Center (ZNZ), University of Zurich and ETH Zurich, Switzerland; Swiss Epilepsy Center, Klinik Lengg, Zurich, Switzerland; Medical Research Council Brain Network Dynamics Unit, Nuffield Department for Clinical Neurosciences, University of Oxford, Oxford, UK; Wellcome Centre for Human Neuroimaging, University College London, London, UK

## Abstract

The entorhinal cortex (EC) supports a coordinate system for spatial memories, organised in a hierarchy along the EC dorso-ventral axis. Recent theories suggest that a similar coordinate system could scaffold non-spatial memories. Here we show that an abstract hierarchical coordinate system supports arbitrary sequence memories in the human medial temporal lobe (MTL). In single-unit recordings from MTL, we find abstract, coordinate-like coding in a simple sequential memory task. In fMRI we find that abstract coordinate representations are arranged hierarchically along the entorhinal cortex, mirroring the anatomical gradient of grid cells in the rodent EC but now for non-spatial sequences. We replicate this finding in an independent cohort of participants. These data suggest that memories are scaffolded on a hierarchical coordinate system aligned to preserved anatomy across domains and species.

---

In fields like mathematics, physics or geography, a coordinate system is a tool constructed to determine the position of an object. To be useful, a coordinate system should be 1) efficient: compressing the position information effectively; and 2) generalisable: appropriate in different situations. Research in the well-studied field of spatial neuroscience has revealed that the brain has constructed a neural coordinate system for knowing one’s position in physical space: grid cells in the entorhinal cortex^1^, which are active on a regular triangular lattice when an animal moves in 2D space. Grid cells are arranged into discrete, factorised modules of different scales and spatial frequencies^2^. Readout from these modules can be combined to efficiently encode current location^3^ and to support inferences such as shortest path to goal^4^. The grid cells code also generalises: it abstracts over the sensory particularities of spatial environments^5–7^. Beyond space, neurons in the entorhinal cortex have also been shown to provide a temporal basis for ongoing experience^8–10^, though whether such codes are organised along an anatomical hierarchy is not known.

Here we set out to ask whether the same coding principles extend beyond physical space to organise arbitrary sequences of experience. Such a coordinate system could serve as a scaffold^11^ for non-spatial memories. For example, the position in the sequence breakfast-lunch-dinner can be thought of as a “stage of day” coordinate and provides a useful reference to encode and retrieve memories. Encoding this sequence with a coordinate entails a generalisable code: all breakfasts share commonalities, enabling abstraction that can aid behaviour in novel situations. Importantly, sequences often have a nested hierarchical structure, with different levels of the hierarchy corresponding to different scales or temporal frequencies. This can allow for an efficient factorial description in a coordinate system with multiple coordinates, akin to grid modules in physical space or to the multiple timescales encoded by entorhinal neurons in time. For example, consider the hierarchy “day stages-days-weeks-months-years”, where each level consists of a sequence from the previous level. You might remember that the first astronauts stepped foot on the moon in 1969, but not that they did so in July. Conversely, you might look back on your vacation in Italy and be confident that you travelled in August, but not remember in which year. A representation of such a hierarchy with a coordinate system can use the structure in sequences to encode them usefully for memory.

We first tested for a representation of a single coordinate of task stage by recording from neurons in the MTL of patients performing a simple sequential memory task. We find cells responding to different task stages, similar to previously observed time cells, but abstracted over sensory input. We then use fMRI in two independent cohorts of healthy volunteers to probe a factorised, multiple coordinate representation of general hierarchical sequences, serving as a scaffold for memory. We aimed to test whether, like the entorhinal codes for space and time, sequence representations are abstract, and organised on an anatomical hierarchy. While our hypotheses are inspired by the properties of spatial grid cells and lateral entorhinal temporal codes, we note that our study did not address whether the proposed sequence coordinate system repurposes these cells or involves different ones. Moreover, while grid cells are organised into discrete modules, our task could in principle be represented either as discrete modules or as a continuous gradient.

## Representation of task stage coordinate in human MTL

Knowing one’s position in the task at hand is particularly useful in tasks with a repeating structure – as is common in lab tasks with same-structured trials. To test if the human MTL encodes a “task stage coordinate” representation in such a task, we recorded cells in hippocampus (92 neurons) and entorhinal cortex (75 neurons) of epilepsy patients performing a memory task with a repeating sequential structure (Fig S1.1 and Fig S1.2). In each trial, patients were shown four pictures in a sequence, and after a delay of 3 seconds were asked to report the sequence position of one of the pictures (Fig 1a). Importantly, while the same 4 pictures were used in every trial, their order was randomised across trials (Fig 1b). Patients performed the task well, with an average accuracy of 93.7% ± 6.3% (SD).

**Fig 1.**
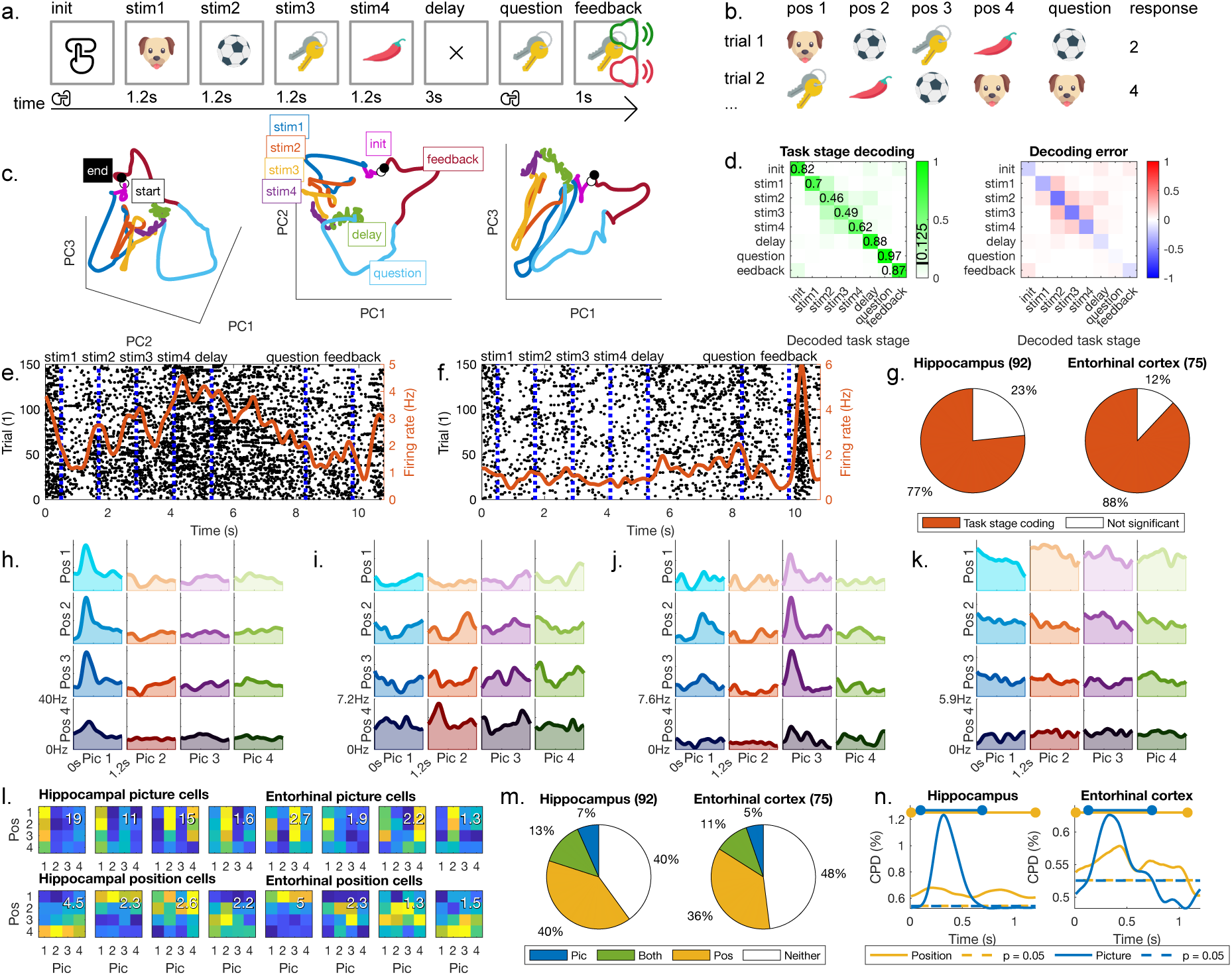
Single unit data. **a.** Participants had to learn the order of 4 pictures presented one after another. After a 3-second delay, we queried the sequence position of one of the pictures. **b.** On each trial, we used the same four pictures but changed the order in which they were displayed. **c.** The averaged population trajectory throughout a full trial of the task in neural activity space shows that different task stages occupy different regions in neural activity space. **d.** Decoding task stage from the population firing rates. Left: average decoded task stage (columns) given true task stage firing rates (rows) across trials, with chance level (1/8) indicated on the color scale. Right: decoding error**. e.** Raster plot across trials for a hippocampal cell showing significant task-stage modulation. The red line is the average firing rate. **f.** Raster plot across trials for an entorhinal cell showing significant task-stage modulation. The red line is the average firing rate. **g.** Proportion of cells showing significant task-stage coding (initiation, stim1–4, delay, question, feedback; one-way ANOVA). **h-l:** Example picture or coordinate cells. We averaged the firing rate of each cell across all trials of a particular picture (column) in a particular position (row). **h.** Hippocampal cell coding for picture 1. **i.** Hippocampal coordinate cell that increases its firing from the first to the last sequence position. **j.** Entorhinal cell coding for picture 3. **k.** Entorhinal coordinate cell that decreases its firing from the first to the last sequence position. **l**. Other examples of picture (top) and position (bottom) specific cells in hippocampus (left) and entorhinal cortex (right) averaged across time within trial stage as a compact variant of the plots in h-k. The maximum average firing rate is indicated by the white number for each cell. **m.** Proportions of picture and position specific cells in hippocampus (left) and entorhinal cortex (right). **n.** Variance explained through time from stimulus presentation by picture and position factors on hippocampus (left) and entorhinal cortex (right).

Visualisation of neural population trajectories strongly suggested that a task stage coordinate was represented, as the activity for different task stages (initiation, stim1, stim2, stim3, stim4, delay, question, feedback) separated in neural space (Fig 1c). Indeed, using a Bayesian decoder, we could decode each task stage from the population activity with high accuracy (Fig 1d, average decoding accuracy of correct task stage 0.73, chance is 0.125). This representation was also present at the single neuron level: A one-way ANOVA revealed that most cells (Hippocampus: 77%, EC: 88%, Fig 1g) in the MTL were tuned to task stage (see example cells in Fig 1e,f). Together, these results indicate that neurons in the MTL overwhelmingly represent different stages of the task – across the population they can code the structure of the task. Note, however, that this representation is not necessarily abstract, as these analyses are partially confounded by external sensory experience (e.g. common visual experience in all trials during the initiation or delay periods, and common auditory experience during feedback) and internal neural processes (e.g. working memory, motivation, motor preparation).

## Coordinate cells in the human MTL encode the position in a sequence abstractly

Next, we asked whether MTL coordinate cells abstracted over the sensory particularities of trials. To test this in a controlled manner, we focused on the presentation periods of the pictures during encoding (stim1-stim4), where our design ensured there were no visual and motor confounds. Such “abstract coordinate cells” would fire, for example, in the third position regardless of the picture shown. An ANOVA with two factors (position and picture) revealed a significant main effect of position in approximately half of the recorded MTL neurons (Fig 1m, Hippocampus: position only 40%, picture only 7%, both 13%; Entorhinal cortex: position only 36%, picture only 5%, both 11%; example cells in Fig 1g-l and Fig S1.3, S1.4; tuning to only position 2 and 3 in Fig S1.5). Interestingly, while the sensory picture code was (expectedly^9^) present transiently after picture presentation onset (blue line in Fig 1n, permutation p<0.05 family-wise error corrected through time: hippocampus: 0.1-0.69s, EC: 0.13-0.74s), the abstract position code was present before and after the presentation of the stimulus (yellow line in Fig 1n, hippocampus: 0.01-1.2s, EC 0.01-1.08s). In summary, these results indicate that in this simple sequential memory task, the human MTL encodes an abstract coordinate representation, independent of sensory stimuli.

The hierarchical structure of sequences is used behaviourally as a scaffold for memory

Our next goal was to investigate whether the MTL’s sequence coordinate system would extend to memory tasks with hierarchical sequential structures, where an efficient representation should comprise multiple coordinates – one for each level of the hierarchy. Indeed, the neural coordinate system for space (grid cells) is organised in exactly such a hierarchical form – grid cell modules (Fig 2a), which compress the spatial code in a factorised manner (Fig 2b). Critically, these modules are anatomically organised on a ventral-dorsal gradient in the rodent brain, from large scale (low frequency) to small scale (high frequency) grid modules^2^ (Fig 2d top). Whether non-spatial sequences are organised along a similar gradient — by grid cells or other entorhinal cell types — is unknown. We hypothesised that hierarchical sequences used for memory will be represented in the human EC with a coordinate system organised on a similar anatomical gradient (Fig 2c) - whether or not this gradient relies on grid cells themselves. Such a finding would resonate with recent theoretical suggestions that grid cells are a compact abstraction of 2D sequences, and that similar principles could apply to sequences drawn from other structural forms^7,12–15^ (though potentially in different cells than grid cells)

**Fig 2.**
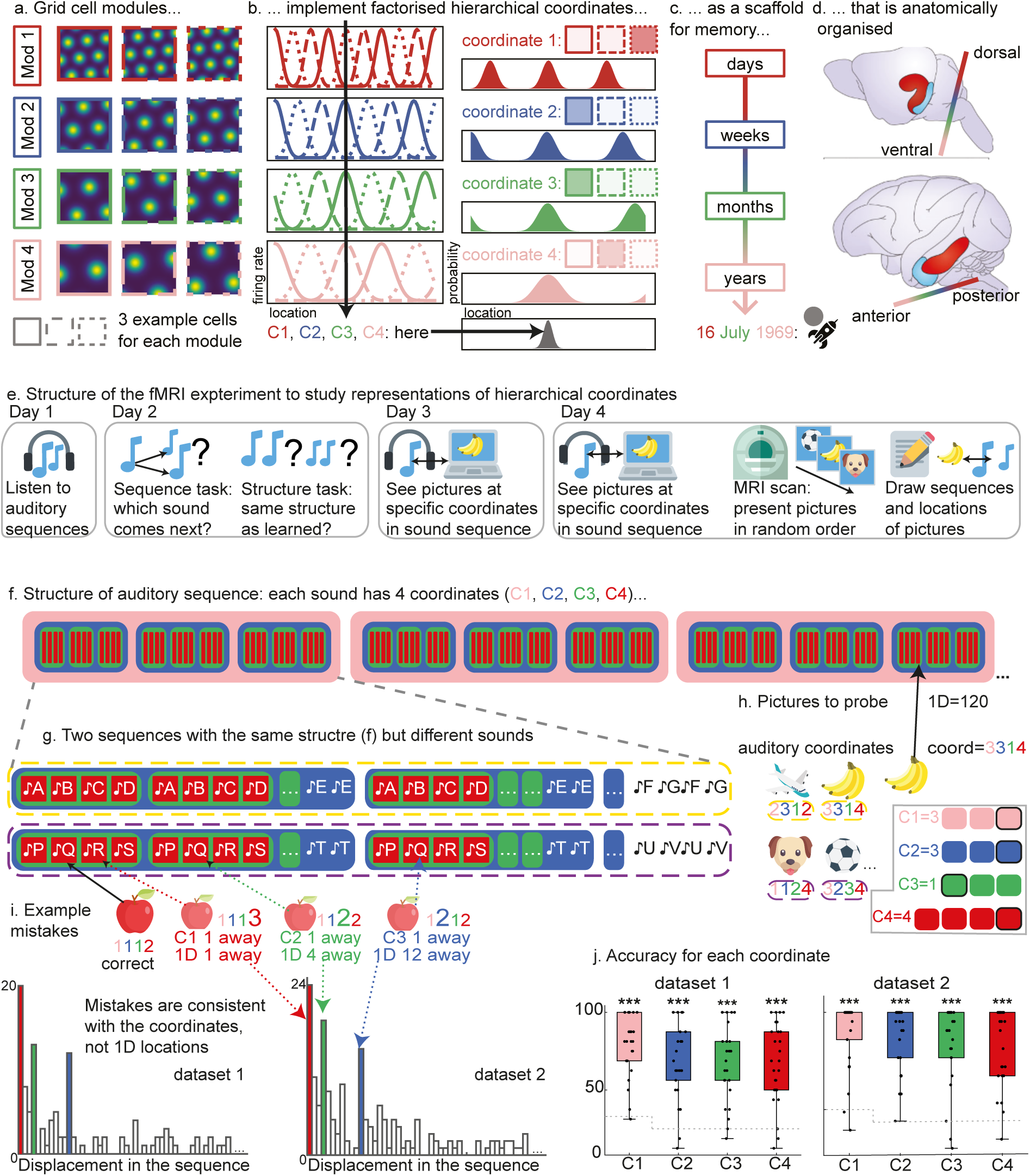
fMRI experimental design and behaviour. **a-d.** Schematic of logic for anatomical hypothesis: coordinates systems organised anatomically by scale in EC**. a-b.** Grid cells are a coordinate system for space. **a.** Simulated example grid rate maps. Grid cells in rodent entorhinal cortex group together in anatomically separable modules of different spatial scales (rows). Cells within a module exhibit different shifts or ‘phases’ (columns, in differently dashed boxes). Adapted from Stemmler et al., 2015^3^. **b**. Together, these grid cell modules implement a factorised hierarchical coordinate system, here illustrated in 1D for clarity. Given the tuning curves of each grid cell in a module (left, different dashes), a particular firing pattern across cells (e.g. cell 3 in module 1 is active while cells 1 and 2 are silent) specifies a particular (module scale dependent) distribution over locations (right) - the readout of the coordinate. Combining the multi-scale coordinates across modules yields a hierarchical code for one specific location (grey, bottom). **c**. We hypothesise that similar factorised hierarchical codes in the human brain provide a scaffold for memories. For example, the hierarchical representation ‘16 July 1969’ corresponds to the memory of the first man on the moon. **d**. For physical space, this gradient of coordinate representations from small to large scales is organised anatomically in rodent entorhinal cortex along the dorsal (small scale grid cells) to ventral (large scale grid cells) axis. In humans, this axis has rotated into the posterior-to-anterior direction. **e**. We test this hypothesis in a four-day neuroimaging experiment. Participants learn auditory sequences (day 1-2) with specific locations tagged by pictures (day 3). We show the same pictures in random order in the MRI scanner on day 4, without any auditory stimuli. After the scan, we ask participants to draw the sequences and place the pictures at their associated locations (an example drawing in Fig S2.2) **f.** Hierarchical structure of the auditory sequences (full sequences in Fig S2.1). **g**. Participants learned two sequences with identical structures but different sounds. **h**. Example pictures and their specific sequence locations that participants learned on the third day. The 1D position of every picture in the auditory sequence can be pinpointed using four coordinates. In the fMRI scanner we showed pictures in a random order without sounds, with the goal of reactivating the representations of the corresponding auditory coordinates. **i**. Distance between incorrectly placed pictures in the post-scan drawings of day 4 and their correct locations. The three strongest peaks in the histogram correspond to incorrectly identifying the coordinate in one level while correctly identifying all other coordinates. **j.** The average accuracy of the post-scan drawings across all pictures for each hierarchical coordinate.

Due to the tilt of the human MTL compared to the rodent’s, the human EC gradient should extend along the anterior-posterior (A-P) rather than ventro-dorsal axis^16^ (Fig 2d bottom). Because obtaining enough cells along the A-P axis in human patients is impractical, we used fMRI. Although this study aimed to measure the organisation of memory, we decided not to rely on real remembered episodes of our participants, as these are easily distorted by e.g. emotional valence or recency biases. Instead, we designed a highly controlled task that induces a particular memory structure to afford precise representational hypotheses. We collected two independent datasets (26 and 22 participants, obtained years apart), and replicated all results across the two datasets.

We designed a task to look for hierarchical organisation of sequence coordinates using auditory sequences. This entailed teaching participants two 131-tones-long auditory sequences by repeated listening and testing of the sequences for 2 days (Fig 2e,f,g). The sequences shared the same nested hierarchical structure, with 4 levels – each level containing 3 or 4 locations (Fig 2f,g and Fig S2.1, Methods). Any specific (flat, one-dimensional) location in this sequence can therefore be pinpointed by 4 coordinates, where here a coordinate is a location within a level of the hierarchy (Fig 2h, right). We hypothesized that these coordinates would be represented along an anatomical gradient in the entorhinal cortex with coarsest level’s coordinate anterior and finest level’s coordinate posterior (Fig 2d).

However, to be certain that we could measure the coordinate representations without biases such as temporal autocorrelation, we needed to be able to present the coordinates in a random order. We therefore could not present the actual auditory sequence in the scanner. Instead, we tagged 8 locations in each sequence with visual images (Fig 2h). To do this, on the third day of auditory sequence learning, participants saw 8 images on a computer monitor at specific sequence locations during 25 repeats of the sequence (Fig 2e,h). This procedure therefore endowed 16 images (8-per-sequence) with 4 sequence coordinates each (one-per-level). On the fourth day, participants heard 10 more repeats of the sequence with the paired images as a reminder, and then underwent fMRI scanning where they viewed the images in a random order, without any auditory tones (Fig 2e). Crucially this meant any structure in the resulting fMRI activity could only be a result of the underlying coordinate representation, and not the statistics of the presentation order. To maintain attention, participants were instructed to respond when the shape of the image was distorted (6% distractor trials). Subsequent analyses look for the neural representations of the *images’ coordinates* in entorhinal cortex.

After the scan, participants drew the sequences, placing the pictures in their correct positions (see Fig S2.2 for an example). Accuracy was significantly above chance for all levels (Fig 2j, t-test against chance: all levels, both datasets p<0.001). The displacement errors were consistent with a hierarchical rather than a flat representation of the sequence: the strongest peaks correspond to mistaking one coordinate of a picture while correctly placing all other coordinates (Fig 2i). Indeed, the distance in hierarchical coordinate space was a better predictor of errors across participants than linear sequence distance (multiple regression, p<0.001 dataset 1, p=0.02 dataset 2).

## A coordinate system for hierarchical sequences is organised on an anterior-posterior gradient in the human entorhinal cortex

To test our hypothesis of an anterior-posterior EC gradient from low to high frequencies, we first needed to quantify the coordinate representation at each of the 4 levels around each EC voxel. Using representational similarity analysis^17,18^ (RSA), we built a hypothesis similarity matrix for each level, which assumed that nearby coordinates in that level would have more similar neural representations. This led to 4 hypothesis matrices (Fig 3b). For each level, the matrix was 16×16, with each element being the predicted similarity between a pair of images’ coordinates at that level. We used these as regressors in a multivariate RSA searchlight analysis^19^. The data matrix, at each voxel, was the correlation matrix between voxel responses in a local searchlight between the 16 images, resulting in a 16×16 data matrix per-voxel. We ran a multiple regression resulting in 4 t-statistic maps in each dataset – one per level (Fig 3c, Methods).

**Fig 3.**
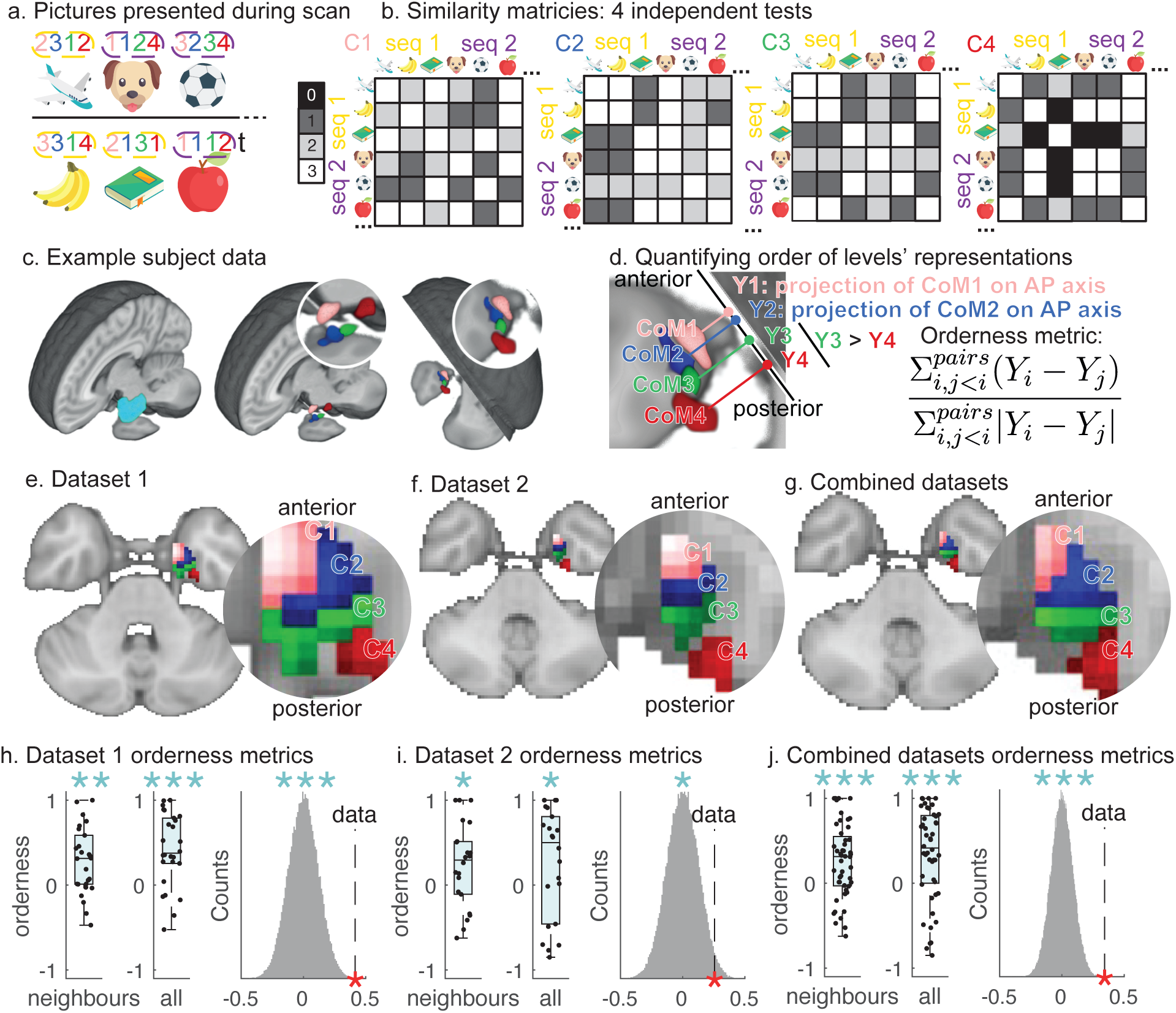
Coordinate representations are anatomically organised. **a.** We present pictures associated to specific coordinates in random order in the MRI scanner. **b.** To find coordinate representations, we specify hypothesised similarities between pictures with respect to each level of the hierarchy. For example, the dog and the book have high similarity for coordinate C2 (blue: both 1), but low similarity for coordinate C4 (red: 4 versus 1). **c.** We use RSA to detect within entorhinal cortex (left) where neural similarities match the hypothesised similarities (example participant, middle), along the anterior-posterior axis (right). **d**. To quantify the A-P order of coordinate representations, we calculate an “orderness metric” from differences between projections (Yi) of each coordinate representation’s centre of mass (CoMi) on the anterior-posterior axis. If the gradient is present, the differences are expected to be positive, because we always subtract the lower from the higher level, and the projection value increases in anterior direction. **e**-**g**. Visualisation of group averaged coordinate representations in dataset 1 (e), dataset 2 (f), and both combined (g). **h**-**j**. Orderness metric for neighbouring levels’ pairs only (left), including all pairs of levels (middle), and averaged across participants against a null distribution from shuffled level labels (right) in dataset 1 (h), dataset 2 (i), and both combined (j).

No voxels survived whole brain comparison for any level (perhaps due to the passive nature of the task, or the extremely small areas of cortex hypothesized to respond to individual levels). Nevertheless, we were able to look for the relative positioning of the different levels’ coordinate effects within entorhinal cortex, and to replicate these relative positions in an independent dataset. Indeed, visual inspection of both independent datasets suggested that for the left EC, these effects were ordered on the A-P axis from low to high frequencies (visualisation Fig S3.1a), as hypothesised (Fig 3c: example participant, Fig 3e: dataset 1, Fig 3f: dataset 2, Fig 3g: both datasets together).

To statistically test for the gradient, we first extracted the centre of mass (CoM) of each level’s map (masked within each hemisphere’s EC), and projected it on the anterior-posterior (A-P) axis, yielding a single A-P location per level in each participant (Y_i_ in Fig 3d, left). We aimed to test whether the order of these projections matched the hypothesised order, in both datasets. We quantified order by calculating an “orderness” metric from distances between ordered pairs of A-P locations (Fig 3d right). This metric computes the sum of differences between the A-P locations of pairs of levels’ representations (positive if the low frequency level is anterior to the high frequency level, as hypothesised), normalised by the unsigned distance. This metric is more positive if low frequency levels are represented more anteriorly than high frequency levels. Indeed, the orderness metric was positive for different combinations of sequence coordinates in both datasets, which confirmed our hypothesised order. First, we calculated the orderness metric only for the neighbouring levels in the hierarchy (e.g. levels 1-2, 2-3 etc.). Indeed, they were ordered according to our hypothesis (Fig 3h-j left, one-sided t-test: dataset 1 p<0.001, dataset 2 p<0.05, combined p<0.001, Supplementary Table 1). The orderness metric including all pairs of levels was also positive (Fig 3h-j middle, one-sided t-test: dataset 1 p<0.001, dataset 2 p<0.05, combined p<0.001, Supplementary Table 1). This measure was significantly larger than a null distribution created by randomly shuffling level labels (Fig 3h-j right, permutation test: dataset 1 p<0.001, dataset 2 p<0.05, combined p<0.001, Supplementary Table 1). All results above are reported in the left entorhinal cortex; we did not find evidence of a gradient in right entorhinal cortex (Fig S3.2, Supplementary Table 2).

Beyond the replication across two independent datasets, the results were also not sensitive to the specific analysis choices we made. The results held when we used a more conservative EC mask (all p< 0.05, Fig S3.3, Supplementary Table 3), and when using a discrete measure of the A-P order of the levels rather than a continuous one (all permutations p<0.05, Fig S3.4, Supplementary Table 4). Furthermore, the locations of the individual levels’ coordinate representations replicated across datasets. On average, a level’s t-statistics were higher when computed at that level’s centre of mass from the other dataset than when computed at the centre of mass of other levels in the other dataset (p<0.01, Fig S3.1b,c).

## The EC’s sequence coordinate system abstracts over sensory stimuli

So far, we have shown that the EC encodes a sequence coordinate system in an efficient hierarchical fashion, and that this coordinate system is organised anatomically in a similar way to grid cell modules - the EC’s coordinate system for physical space.

Notably, the grid representation is also abstract – another important property of coordinate systems: within a grid module, the covariance structure of grid cells’ activity is maintained across different spatial environments^5^. In other words, the information embedded in grid cells abstracts over sensory details. EC codes for time show a related form of abstraction, being reshaped by the structure of events rather than tracking sensory experience alone^8,9^. To test whether this was also true for our sequence coordinates, we re-did the entire analysis above, but now selecting only the portions of the hypothesis matrices which compared images from *different auditory sequences* (Fig 4b). This means any similarities between the data and the coordinate hypothesis matrices could not be because of sequence-specific reasons. We then followed the same procedure for the analyses as described above (group statistical maps in Fig 3c-e). Despite the reduced power of the analysis (using only half of the hypothesized correlations), all permutation tests were significant (Fig 4f-h right, permutation test: dataset 1 p<0.001, dataset 2 p<0.05, combined p<0.0001, Supplementary Table 6) indicating an abstract generalisable coordinate system.

**Fig 4.**
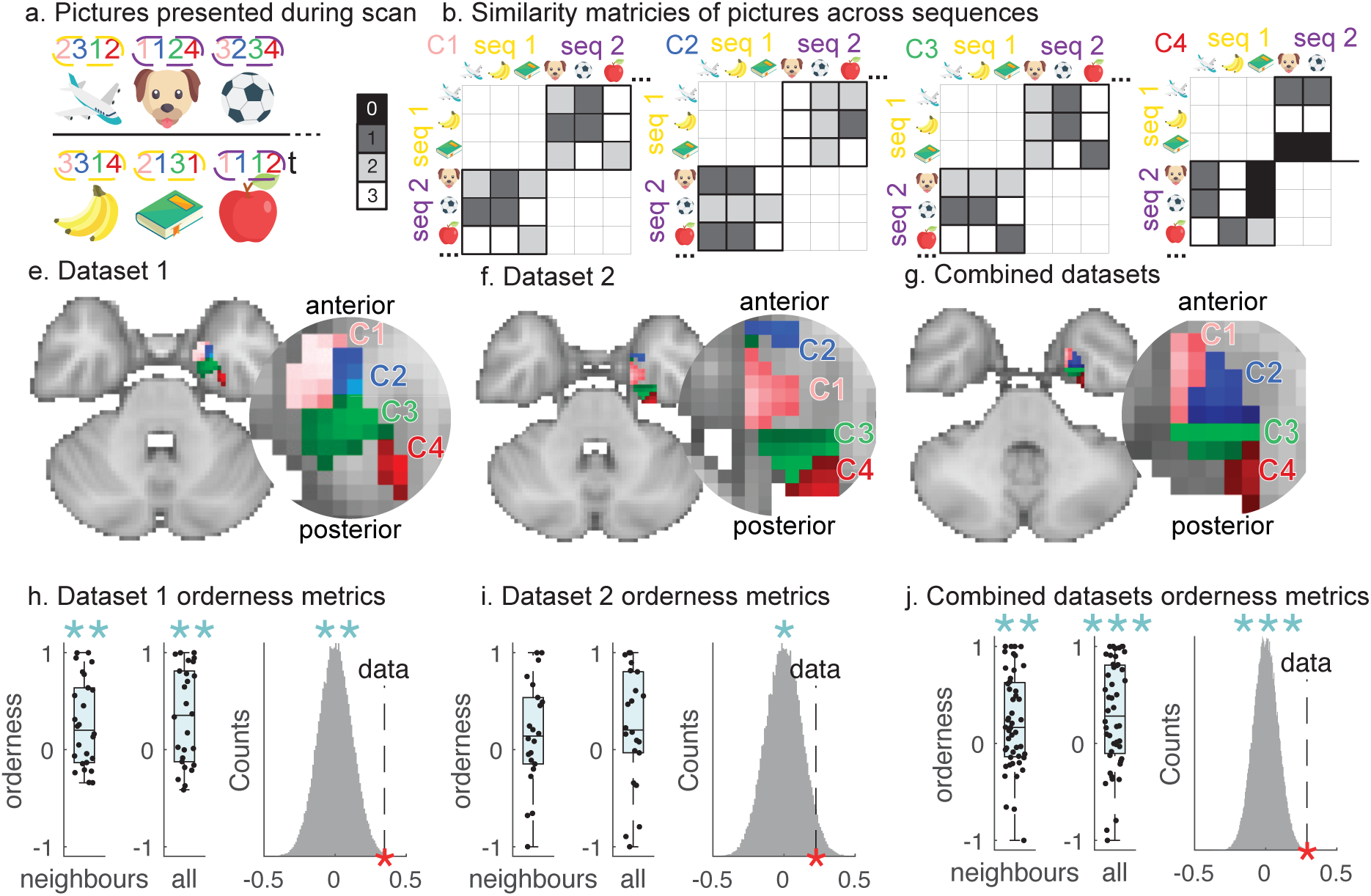
Abstract generalisable coordinate system. **a.** We presented pictures associated to specific coordinates in random order in the MRI scanner. Notice that some were paired with a sequence 1 coordinate during training (yellow dashed line) and others with sequence 2 (purple dashed line). **b.** Across-sequence similarity matrices. To find coordinate representations, we specify hypothesised similarities between pictures with respect to each level of the hierarchy. This time we include only similarities between pictures from different auditory sequences for further analysis. **c-e**. Visualisation of group averaged coordinate representations in dataset 1 (**c**), dataset 2 (**d**), and both combined (**e**). **f**-**h**. Orderness metric for neighbouring levels pairs only (left), including all pairs of levels (middle), and averaged across participants against a null distribution from shuffled levels’ labels (right) in dataset 1 (**f**), dataset 2 (**g**), and both combined (**h**).

## Discussion

Understanding the temporal structure of experiences is a key aspect of learning and memory. Here, we show that the brain encodes this structure with an abstract coordinate system, anatomically organised hierarchically for efficient encoding. Using single-unit data, we first show that in a task with a repeating structure the MTL encodes a stimulus-invariant task-stage coordinate. We then show using fMRI that in a task with a hierarchical structure the EC encodes multiple factorised coordinates which are arranged anatomically on the anterior-posterior EC axis according to their level in the hierarchy, mirroring the organisation of grid cell modules in spatial tasks. Taken together, these results suggest that a coordinate system-like representation might be a general shared representational principle across multiple tasks and domains.

Previous studies have described gradients of temporal and spatial granularity outside of entorhinal cortex, highlighting a broader principle of hierarchical organization in the human brain^20–24^. However, these studies are looking for place-cell-like representations that vary in their resolution. In contrast, our study was designed to look for a factorial hierarchy (like the grid code), that repeats with different periodicities at different levels of the hierarchy. In principle it would be possible to check for other brain regions displaying the same factored hierarchy, although given the nature of the gradient analysis, this would require strong hypotheses about the gradient direction and boundaries in each anatomical location. Indeed, we were unable to find such a hierarchy even within right EC. Unilateral effects are common in fMRI studies of grid-like coding^25–27^. While the reason for our left lateralization remains unclear, the replication across two independent datasets supports the robustness of the effect.

Previous studies have found representations of time in the medial temporal lobe. Time cells that respond to intervals between events exist in rodents^8,9,28–31^ and humans^32^. These include codes that are hierarchical, with ramping responses over multiple timescales, but not abstract, because they are tethered to sensory experience^8^. Others are abstract but not hierarchical, with transient responses that tile a single timescale^28–32^. In our studies, such a representation would account for the data only if the representations of time were restructured according to the structure of events. In the cells, where the structure is linear, this means representations that tile time from the start to the end of each trial. In the fMRI, it would require an even larger restructuring, with time being measured independently for every level in hierarchy, with each level perfectly matching the same level in the auditory sequence. It is already known that temporal codes in the rodent lateral entorhinal cortex are indeed reshaped by simple event structures^8,9^. However, so far this has been described for laplace-like representations that peak at the start and end of an event. Our cells do not have this form, as many peak for intermediate points in the sequence (as do previous coordinate cells for spatial sequences^33,34^).

But our experiences are not distributed uniformly in space and time. They form richly structured sequences. Statistical dependencies in experience, for example sequences of goals in space^26,33,34^ and graph structures embedded in time^10,35–37^, do indeed influence both fMRI signals and cellular responses in the human entorhinal cortex. These responses have been suggested to support the formation of a cognitive map: an internal representation that allows animals to reason about future outcomes. Learning a cognitive map requires understanding the relational structure that underlies experience and representing it in a way that facilitates flexible behaviour. Our study shows that such representations for general sequences beyond space and time can be abstracted to an efficient coordinate that compresses the experiential structure.

Strikingly, this hierarchical compression aligns with the anatomical gradient of the entorhinal cortex in exactly the fashion predicted by the grid cell hierarchy, though our data cannot identify the cellular substrate of this gradient, which could involve grid cells in the medial entorhinal cortex, time and task cells in the lateral entorhinal cortex, a combination of these, or other entorhinal populations. This raises the question of whether the two hierarchies are encoded by the same cells or whether there are separate abstract coordinate systems across domains. It is not possible to know this from our study. Recent theoretical work offers reasons to favour both possibilities^11^. Forming a bespoke coordinate system that captures the experiential statistics allows efficient inferences and predictions about unseen events^7,12,38^ and efficient memory search^39^. By contrast, in similarly structured models, using grid cells themselves as the coordinate scaffold for arbitrary non-spatial sequences provides stable memories that gracefully degrade at capacity limits^11^. Testing between these ideas will likely require studies that measure representations of space and structured non-spatial sequences in the same cells. However, in either case, our data establishes that the anatomical hierarchy in the entorhinal cortex is present in humans and broadens its classical understanding beyond 2-dimensional spatial sequences to arbitrary sequences of experiences.

## Methods

### Intracranial data: participants

Intracranial recordings were acquired from 6 patients with pharmacologically resistant epilepsy in the intensive monitoring unit of the Klinik Lengg in Zurich, Switzerland. The ages of the participants ranged from 29 to 56 years (mean = 38.6, SD = 11.6). As part of their evaluation for neurosurgical epilepsy treatment, all patients had sEEG depth electrodes intracranial electrodes implanted in the medial temporal lobe for diagnostic purposes. During the experiment, no clinical seizures occurred. Informed consent was given by all patients. All participants provided written informed consent for the study, which was approved by the institutional ethics review board (PB 2016-02055). All participants had normal or corrected-to-normal vision.

### Intracranial data: the task

The patients had to memorise a simple four-picture sequence. The same pictures were shown in every trial, but their order was randomised. Each image was shown for 1200 milliseconds at the center of the screen with no interstimulus interval. There was a 3-second delay after the sequence was presented. Following that, one of the pictures was presented, and the patients had to answer where it was in the sequence (position 1, 2, 3, or 4). Before the next trial, auditory feedback indicated whether the given response was correct. Each session included 50 trials, and most patients completed multiple sessions of the task (mean = 3.3, SD = 1.2).

### Intracranial data: recordings setup

The depth electrodes (1.3 mm diameter, 8 contacts of 1.6 mm length, and spacing between contact centres of 5 mm; Ad-Tech, Racine, WI, www.adtechmedical.com) were stereotactically implanted into amygdala, hippocampus, and entorhinal cortex (Fig. S1.1a). In this work, we only focused on the recordings from entorhinal cortex and hippocampus (Fig S1.1b). Each macroelectrode had nine microelectrodes that protruded approximately 4 mm from its tip. The data from the microelectrodes was acquired at sampling frequencies of 32 kHz via the ATLAS recording system (0.5- to 5000-Hz passband, Neuralynx, Bozeman, MT, USA; www.neuralynx.com). The recording setup is described in detail in^40,41^.

Anatomical localisation of implanted electrodes was performed with Lead-DBS, a specialized software for electrode mapping^42^. This process involved co-registration of pre-implantation MRI and post-implantation CT, normalization of pre-implantation MRI to MNI space, subcortical refinement to account for brain shift and manual annotation of electrode contact positions. Additionally, each patient’s pre-implantation MRI was segmented with FreeSurfer^43^, by applying cortical parcellation and subcortical structure segmentation to precisely determine anatomical electrode locations. The automated segmentation was verified, with a clinician’s assessment serving as the final determination of electrode localization. The position of the microwire channels was assigned to the location of the first macro contact.

Spike sorting was performed using the Wave Clus software^44^, followed by manual refinement to ensure data quality. The first multi-unit cluster was excluded to ensure we have only single-unit data. Clusters in which spikes were predominantly observed within noisy segments of the recording were removed, as were those exhibiting non-physiological noise or high proportion of inter-spike intervals (ISI) shorter than 3 ms. Additionally, clusters with highly similar waveforms were merged to prevent over-clustering (Fig S1.2).

### Task structure

To explore whether neurons in the human hippocampal formation encode the structure of a task, we first visualised the averaged population trajectory throughout a full trial of the task in neural activity space. We calculated the firing rate of each unit in 10 ms bins, except during the initiation and question stage. These stages of the task had a variable duration across trials because they only end when the patient responds. To align these stages across trials, we divided them into a fixed number of 50 (initiation) and 150 (question) bins in each trial, so that the bin size may vary from trial to trial. We then averaged the firing rate across trials for each unit so that we ended up with a NxT matrix of activity, where N is the number of cells and T the number of time bins. We did a principal component analysis and projected the data on the first three principal components to get a 3xT matrix that described the trajectory in neural space after dimensionality reduction. We smoothed this trajectory with a gaussian kernel (using matlab’s smoothdata) of 25 bins (250 ms) before plotting (Fig 1c).

Since different task stages occupied different regions in neural activity space, we expected that task stages could be separated by their patterns of activity. To test this, we decoded task stage from the population firing rates using a Bayesian decoder. We calculated tuning curves for task stages by averaging the firing rate within each task stage across all odd trials, and then calculated the decoded posterior probability of each stage given the averaged firing rate within that stage for all even trials. We then repeated this procedure the other way around, decoding the task stage from the average of odd trials using tuning curves from averaging even trials. We averaged these two repeats, then plotted the resulting probability of decoding each task stage given the firing rates of each task stage (Fig 1d left). To calculate the decoding error, we subtracted the identity matrix from the decoding probabilities (Fig 1d right). We also plotted raster plots with overlaid average firing rate (after gaussian smoothing with a 50-bin kernel) across trials for two example cells that exhibit strong task structure coding (Fig 1e, f).

### Sequence position

The stimulus presentation stages of the task provided the cleanest separation of structure and sensory input, because the stimuli were randomised across positions across trials. In each trial, we presented the same four stimuli, but in a random order, so that responses to stimulus identity and sequence position could be identified separately. For each stimulus and each sequence position, we collected all occurrences where that stimulus appeared in that position. We averaged firing rates across these occurrences for each 10 ms time bin for each neuron. We then smoothed the firing rates with a gaussian kernel (using matlab’s smoothdata) of 30 bins (300 ms) to obtain a neural response through time for the given stimulus in the given position for each neuron (Fig 1h-k). Additionally, for a more compact representation of the same data, we first averaged the firing rate through time in each stimulus-in-position occurrence, and then averaged across occurrences, to obtain a 4×4 (position by stimulus) matrix of average firing rates (Fig 1l).

We then asked whether either of these variables, i.e. stimulus identity and sequence position, significantly modulated the firing rate of individual neurons. To do so, we averaged each unit’s firing rate across time for each stimulus presentation in each trial, to get a 4*K-dimensional vector with K the number of trials and ran a two-factor Anova (type III sum of squares) where one factor is the stimulus identity and the other the sequence position. We then counted the number of units that come out significant for either or both (Fig 1m).

Finally, we calculated the coefficient of partial determination for stimulus identity and sequence position across the neural population. We started from the 4*K-dimensional vector of stimulus responses at each timepoint for each unit. We then fitted generalised linear models that modelled stimulus identity and sequence position responses with three regressors each. All three stimulus regressors contained a -1 for when stimulus 1 was presented, and each a +1 for stimulus 2, 3, and 4 respectively; similarly, all position regressors contained a -1 for position 1, and each a +1 for position 2, 3, and 4 respectively. We calculated the sum of squared residuals both for the full model including stimulus and position regressors, and a partial model without the regressors for one of those. The coefficient of partial determination was given by the difference between the sum of squared residuals of the partial and the full model, divided by those of the partial model. After calculating the coefficient of partial determination for each timepoint for each unit, we averaged across units to end up with one curve for stimulus identity and one for sequence position, then smoothed with a gaussian kernel of 30 bins (Fig 1n). In order to assess significance, we repeated the exact same procedure but shuffled the labels for stimulus position and identity and built a distribution of the maximum coefficient of partial determination through time for each permutation. The significance level was given by the 95% percentile of that distribution.

### fMRI experiments: participants

We carried out two fMRI experiments with the same task but two independent cohorts of participants. 30 healthy participants participated in each study (dataset 1: 30 participants aged 20–30, mean = 23.7, SD = 2.90); dataset 2: 30 participants aged 19–35, mean = 23.8, SD = 3.6). We excluded the participants that did not reach the training criteria on the second day of the experiment (see below for an explanation of the day 2 training criteria; 2 participants were excluded from dataset 1 and 4 participants from dataset 2). We then analysed data only from the participants who drew the structure of auditory sequences correctly on day 4 (see below for an explanation of the day 4 testing criteria; 2 participants were excluded from dataset 1 and 4 participants from dataset 2). Hence, all the analysis presented is based on data from 26 participants in dataset 1 and 22 participants in dataset 2. Monetary compensation was £15 per hour of fMRI scan, with 20p for each catch trial in the scanner and £10 per hour for the behavioural tasks. The study was approved by the local ethics committee (ethics number R43594/RE001). Each individual signed an informed consent form prior to the start of the experiment.

### fMRI experiments: auditory sequence design and behavioural training

To mimic the hierarchical structure of real-life experience of time (meals make up days, days make up weeks, and weeks make up months), we designed two auditory sequences (Fig 2f,g and Fig S2.1) with an analogous four-level hierarchical structure. Crucially, each sequence’s sounds were different (Fig 2g and Fig S2.1). This allowed us to ask whether the neural representation of the structure generalised across different sensory stimuli. Both sequences were 65.5 seconds long in total. Each sound lasted for half a second. We familiarised our participants extensively with the auditory sequences by behavioural training over multiple days (Fig 2e).

Day 1. The participants received audio recordings of both sequences on the first day. To ensure that participants paid attention to the recordings (as training was conducted at home on this day), we asked them to count the number of times a sequence fragment appeared in each recording. The behavioural training took 30 minutes on average, during which each participant heard each sequence 28 times.

Day 2. On the second day, we trained the participants until they learned the auditory sequences very well. The behavioural training consisted of two different tasks, and the training criterion was 100% accuracy on both tasks. The participants’ first task required memorising the order of individual sounds in each sequence. Participants were presented with a sound and asked to determine which of three subsequent sounds was the next one in the sequence. Participants were given three seconds to respond, and each sound was presented for half a second. The second task emphasised on learning the sequence structure. Participants were asked to determine whether a test sequence was identical to the actual sequence (’correct’) or whether it had been altered (’wrong’). There were six test sequences in total, three of which were identical to the original sequence and three altered in one of the levels. Alternations in a level were created by changing the number of repetitions in that level compared to the original sequence. Each training block consisted of participants hearing the true sequence three times and then performing both tasks.

Day 3. On the third day, we added pictures to the audio sequences. Each auditory sequence had eight pictures, each of which was presented in a unique location in the sequence at the same time as the sound it was paired with (0.5 s). We excluded sounds that indicated boundaries (sound E,F,G/T,U,V in Fig 2g) for pairing with pictures. Picture locations were randomly distributed across the participants. We used 4 objects in 4 different orientations in the first experiment to make 16 pictures, and 16 different objects in the second experiment. This manipulation of the orientations of pictures was designed to ask a question in the first dataset which we do not investigate in this manuscript. While passively watching and listening to the audio sequence, participants were asked to remember the location of each visual stimulus in the sequence. Each sequence was played for the participants 25 times.

Day 4. Similarly to day 3, participants were presented with each of the two sequences ten times prior to the scan task to refresh their memories of them. We then acquired fMRI measurements while projecting the pictures at the center of a screen in a pseudorandomized order, without sounds. Thanks to this order randomisation in the scanner we avoid autocorrelation confounds while we measure the coordinate representation associated with the picture on the screen. As a cover task, we instructed participants to press a button whenever a picture was stretched horizontally or vertically (the distractor picture). Every picture was presented twenty times in its original form and once as a distractor image with different sizes. Each participant completed 4 blocks of the task, with 336 trials in each block. The pictures were shown for 0.8 seconds on each trial. Each visual stimulus was presented with a jittered inter-trial interval (ITI) selected from a truncated gamma distribution with a mean of 2.5 s.

After the fMRI scan task, we tested how well the participants remembered the location of each visual stimulus in the audio sequence. First, participants were asked to draw the structure of sequences 1 and 2 with pen and paper. They were then asked to locate the position of each of the 16 different pictures in their drawing of the sequences (for an example drawing see Supplementary Fig 2.2).

### fMRI experiments: behavioural analysis

Custom MATLAB scripts (Mathworks, Version R2018/b) were used for all statistical analysis. To determine the accuracy of each picture placement in each level by each participant, we examined the participants’ post-scan drawings. We calculated their accuracy by comparing the positions of each of the 16 pictures in the participant’s drawing to the coordinate of those pictures in each hierarchical level of the true sequence. The percentage accuracy in each level was compared to chance in a one-tailed t-test to determine whether participants remembered the locations of the pictures in the auditory sequences. The chance levels for levels 1-3 were 33% and 25% for level 4.

Then, we analysed the participants’ errors. We calculated the distance from the correct location for each incorrectly placed picture. These error distances are an indication of whether the participants exploited the hierarchical sequence structure to remember picture positions, through an increased likelihood of hierarchical errors over just linear proximity. In the days-weeks-months analogy, this is equivalent to misremembering that you had dinner with friends on Wednesday, a week later, instead of the immediately following Saturday—despite the latter being closer in time. We counted the number of errors for each error distance and regressed those on the linear error distance and the hierarchical error distance in a multiple regression. We defined the hierarchical error distance as the Euclidian distance between the true four-dimensional hierarchical coordinate and the erroneous four-dimensional coordinate. This regression thus reported to what extent the linear distance and the hierarchical distance explain the number of errors across participants. To determine if the hierarchical distance predicted errors better than the linear distance, we did a one-sided t-test on the contrast of parameter estimates between the two.

### fMRI experiments: fMRI data acquisition

We presented visual stimuli on a screen in the scanner. MRI data were collected from a 64-channel head coil on a 3-Tesla Siemens Magnetom Prisma MRI scanner. A multi-band sequence was used with a multi-band factor of 3 (MB3). Slicing was done with an in-plane resolution of 2×2mm with no breaks and a slice thickness of 2mm each. The slices were acquired in interleaved order, with an echo time (TE) of 20 ms and a repetition time (TR) of 1.235 s.

The data were collected in four separate blocks per participant. The number of acquired measurements varied per block, with an average of 867 volumes per block.

A field map of dual-echo time with whole brain coverage was collected with a voxel size of 2 × 2 × 2 mm3 (1st TE: 4.92 ms, 2nd TE: 7.38 ms). T1-weighted structural images were collected for each participant for co-registration with echo-planar images (EPI). Each T1-weighted image is constructed of 192 axial slices of 1mm each, in-plane resolution = 1×1 mm2, with the following parameters: field of view (FoV) = 192 mm, TE= 3.97ms and TR = 1.90s.

### fMRI experiments: fMRI data preprocessing

Pre-processing was performed using tools from the fMRI Expert Analysis Tool (FEAT), part of FMRIB’s Software Library (FSL^45^). Data for each of the 8 blocks was preprocessed separately. Each block was aligned to the first, pre-saturated image using the motion-correction tool MCFLIRT^46^. Brain extraction was performed using the automated brain extraction tool BET^47^. All data were high pass temporally filtered with a cut-off of 100 s. Registration of EPI images to high-resolution structural images and to standard (MNI) space was performed using FMRIB’s Linear and Non-Linear Registration Tool (FLIRT^46^ and FNIRT^48^), respectively. The registration transformations were then used to map each block’s EPI data to the native structural space. No spatial smoothing was performed during pre-processing.

### fMRI experiments: RSA

Using representational similarity analysis (RSA^17^), we identified how the brain represented the coordinates of the four levels of auditory sequences. We created four independent hypothesis representational dissimilarity matrices (RDMs) by calculating the distances between all the pictures in all four levels, as each image had a coordinate in all four levels of the hierarchy.

The RSA pipeline was built on the RSA toolbox^18^. The preprocessed data were first transformed from functional to MNI standard space. By defining a mask from the Harvard-Oxford Subcortical Structural Atlas in fsleyes, only cerebral cortical and hippocampal voxels were included. Within this mask, we defined 100-voxel searchlights for each voxel. Then, we calculated the average stimulus responses in each searchlight by fitting a general linear model (GLM) to the BOLD data of every voxel within the searchlight. This GLM modelled every one of the four task blocks separately. Each block contained onset regressors for each of the 16 images, as well as separate regressors for catch trials and button presses. We also added motion outlier frames and the average signal in a CSF region of interest as nuisance regressors. Except for nuisance regressors, SPM12 (Wellcome Trust Centre for Neuroimaging, https://www.fil.ion.ucl.uk/spm) convolves regressors with a hemodynamic response function and fits the GLM to produce a searchlight activation pattern: the average response to a stimulus for each voxel in the searchlight. Given the activation pattern for each stimulus in each block, we calculated across-block RDMs X^AB^_ij_ = 1 – mean(corr(x_i_^A^, x_j_^B^), corr(x_j_^A^, x_i_^B^)) where x_y_^Z^ is the activation pattern x for stimulus y in block Z; the mean ensures this dissimilarity matrix is symmetric. We then averaged the across-block RDM X^AB^ across all combinations of blocks where A≠B. By excluding any similarities between pictures within the same block, we avoid biased statistics due to regressor correlations^49^. As a result, we end up with one 16×16 data RDM, containing dissimilarities between each pair of the 16 pictures. As the last step in the within-participant RSA pipeline, we used multiple regression where we regressed the upper triangle of the data RDM on the upper triangle of each of the four hypothesis RDMs (one for each coordinate) in every searchlight. Alternatively, the four hypothesis RDMs could be regressed separately; this produces similar results (Fig S3.5, Supplementary Table 5). The resulting parameter maps were then smoothed with a 5 mm full width at half maximum kernel.

The result of the above-described analyses is a set of brain maps containing regression coefficients in each voxel for each participant. Then, using fMRI Expert Analysis Tool (FEAT)^45^, we calculate statistical maps at the group level to assess the reliability of these results across the entire participant population^45^. The GLM at the group level contained a mean effect regressor.

### fMRI experiments: orderness metric

Next, we asked if the representations for each level were arranged in a gradient along the anterior-posterior dimension of each participant’s entorhinal cortex. To do this, we first identified the location of the centre of mass (CoM) of each level’s effect within the left entorhinal cortex (parallel analyses in right entorhinal cortex yielded a null result: Fig S3.2, Table 2), defined by the Juelich atlas^50^ anatomical mask thresholded at 50% (results for a stricter 75% mask threshold are reported in Fig S3.3, Supplementary Table 3). We extracted each coordinate CoM’s location along the A-P axis, yielding a single number for coordinate i referred to as Y_i_. Then, we determined if the Y values were consistent with an ordered distribution, more so than chance. To quantify to what extent the A-P locations were distributed along the hypothesised order, we defined an ‘orderness metric’.

The orderness metric was defined as follows: 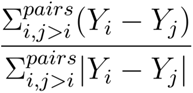. Importantly, for each pair of coordinates included in this measure, we always subtracted Y for the coordinate with the *higher* index from the Y for the coordinate with the *lower* index. To be consistent with our hypothesised order, this difference should be positive, because 1) we have defined Y to increase in the anterior direction of the A-P axis, and 2) the lower-index (lower frequency) coordinate is expected to be more anterior than the higher-index (higher frequency) coordinate. Because we divided by the absolute value of the difference, this metric was normalised between –1 and 1 where 1 indicated perfect ordering, so Y_1_ Y_2_ Y_3_ Y_4_ in A-P order, and –1 opposite ordering, so Y_4_ Y_3_ Y_2_ Y_1_ in A-P order. We separated contributions to the orderness metric from neighbour coordinates versus all coordinates by calculating it twice: once with only neighbouring pairs, i.e. ([1,2], [2,3], [3,4]), and once with all pairs, i.e. ([1,2], [1,3], [1,4], [2,3], [2,4], [3,4]). Finally, we also calculated a discrete version of the orderness metric, that incorporates the sign of the difference rather than the continuous difference value: (results in Fig S3.4, Supplementary Table 4).

We calculated the orderness metric for each participant, both including only neighbouring pairs and including all pairs of coordinates. To evaluate whether the distribution of coordinate representations was significantly ordered as hypothesised, we tested whether the orderness metric was significantly positive across participants. First, we evaluated the significance of a one-tailed t-test against zero of orderness metrics across the population. Then, we constructed a null distribution of population orderness metrics by randomly permuting the coordinate labels for each participant, recalculating the orderness metric for the permuted labels, and averaging the orderness metric across the population. We produced population orderness metrics for 100000 of these permutations. We calculated the permutation test p-value as the fraction of permutations with a larger orderness metric than the population-averaged orderness metric with unpermuted coordinate labels.

### fMRI experiments: ROI analysis

We next tested if the centres of mass (CoM) of each level’s coordinate representation were in the same anatomical locations in entorhinal cortex across the two datasets. We first used the group maps to identify the CoM of each level in the first dataset. We then calculated the effect of each level for each participant in the second dataset at the CoMs defined from the first dataset. We then repeated the procedure but now defined the CoM from the second dataset and tested the effects in the first dataset. We average these two 4×4 matrices together to get a 4×4 matrix E_ij_ containing the effect of level i in dataset 1 at the CoM of level j in dataset 2, averaged with the effect of level i in dataset 2 at the CoM of level j in dataset 1, for each participant. We then compared whether congruent effects were stronger than incongruent ones (e.g., the effect of the third level at the peak of the second level defined in the different dataset vs. the peak of the second level). To do so, we performed a one-tailed paired t-test across participants between the averages of the diagonal elements (congruent effects) and the averages of the off-diagonal elements (incongruent effects).

### fMRI experiments: generalisation analysis

Finally, we tested if the coordinate representations generalised across the two sequences, which had the same structure but different sounds. We again calculated hypothesis RDMs but now the distances for each level were defined only based on the distances between pictures in different sequences (Fig 4b). We then performed the same orderness metric analysis as described above. We also performed the same analysis for the hypothesis RDMs based only on the distances between the pictures within the same sequences.

## Acknowledgements

We thank the participants for their engagement with this difficult task. We thank Salman E Qasim, Joshua Jacobs, Timothy H Muller and Yunzhe Liu for their insightful comments and practical help. We also thank Emile Radyte and Tess Van Stekelenburg for help in the analysis process.

A.S. acknowledges a Wellcome Trust studentship and NIH funding. T.E.J.B. is supported by a Wellcome Principal Research Fellowship (219525/Z/19/Z), a Wellcome Collaborator award (214314/Z/18/Z), and by the Jean Francois and Marie-Laure de Clermont Tonerre Foundation. The Wellcome Centre for Integrative Neuroimaging and Wellcome Centre for Human Neuroimaging are each supported by core funding from the Wellcome Trust (203139/Z/16/Z, 203147/Z/16/Z). The Sainsbury-Wellcome centre is supported by core funding from the Wellcome Trust (219627/Z/19/Z) and the Gatsby Charitable Foundation (GAT3755) and the Gatsby Initiative for Brain Development and Psychiatry (GAT3955).

## Author contribution

Conceptualization and experimental design: A.S., H.B., T.E.J.B. Single unit data collection: A.S., E.M.S., J.S., D.L., L.I. FMRI data collection: A.S. and E.M.S. Data analysis: A.S., J.J.W.B., A.K., H.C.B., T.E.J.B. Manuscript writing: A.S., J.J.W.B., A.B.B, T.E.J.B. Manuscript editing: A.S., J.J.W.B., A.B.B., J.S., D.L., L.I, A.K., E.M.S., H.C.B., T.E.J.B.

## Supplementary Material

**Supplementary figure 1.1.**
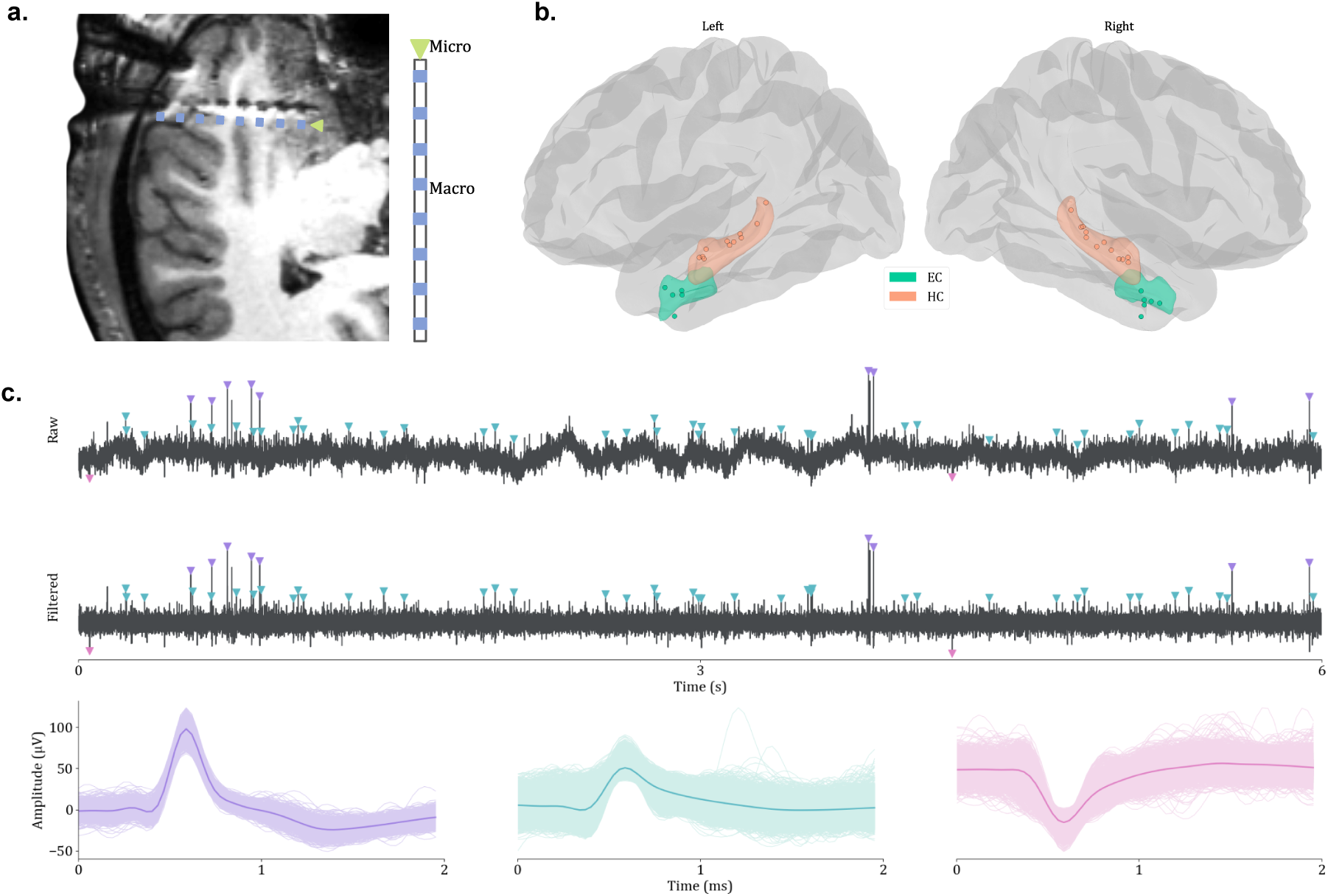
a. Combinatorial electrode. Right: schematic showing macro- (blue) and microelectrode (green) contacts along the depth electrode. Left: example MRI from a patient illustrates electrode placement, with visible macro- and microcontacts, with an overlaid schematic of the electrode positioned slightly below its actual location in the MRI. b. Electrode localization in MNI space, showing electrode positions in the Freesurfer average brain model, with the entorhinal cortex (EC, green) and hippocampus (HC, orange) highlighted. Electrodes are color-coded according to their respective anatomical regions. c. Spike sorting example, showing raw recordings (top) and recordings filtered in 300-3000 Hz range (middle) from a single microelectrode channel. Detected spikes are marked and color-coded corresponding to the three identified neurons (bottom). Individual spike waveforms (lighter color) and averaged waveforms (darker color) are displayed for three identified neurons.

**Supplementary figure 1.2.**
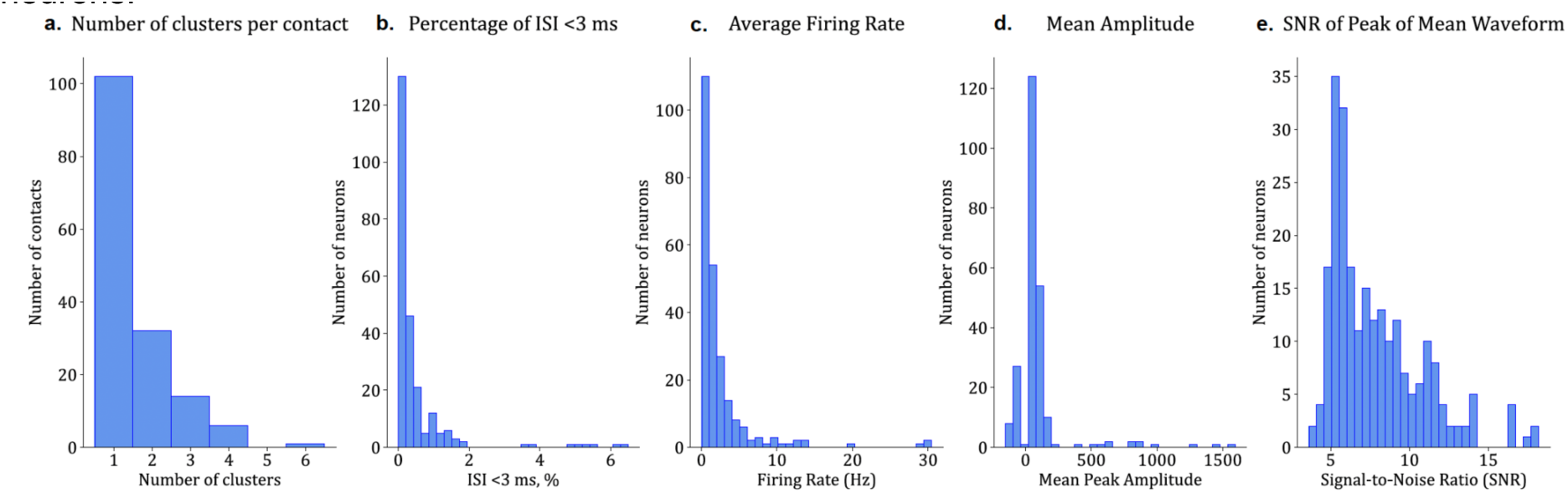
Spike Sorting Quality Assessment. Summary of key metrics used to evaluate the reliability of spike sorting: **a.** the number of neuronal units identified per electrode contact, **b.** the percentage of inter-spike intervals (ISI) shorter than 3 ms, **c.** the average firing rate in Hz **d.** the peak amplitude (µV) of detected spikes, calculated as the mean across waveform peaks for identified neurons, and **e.** the signal-to-noise ratio (SNR) of the peak of the mean waveform, defined as the ratio of the peak amplitude of the neuron’s mean spike waveform to the background noise level (calculated as the peak of the averaged waveform divided by the mean of the standard deviations of individual spike waveforms).

**Supplementary Table 1.**
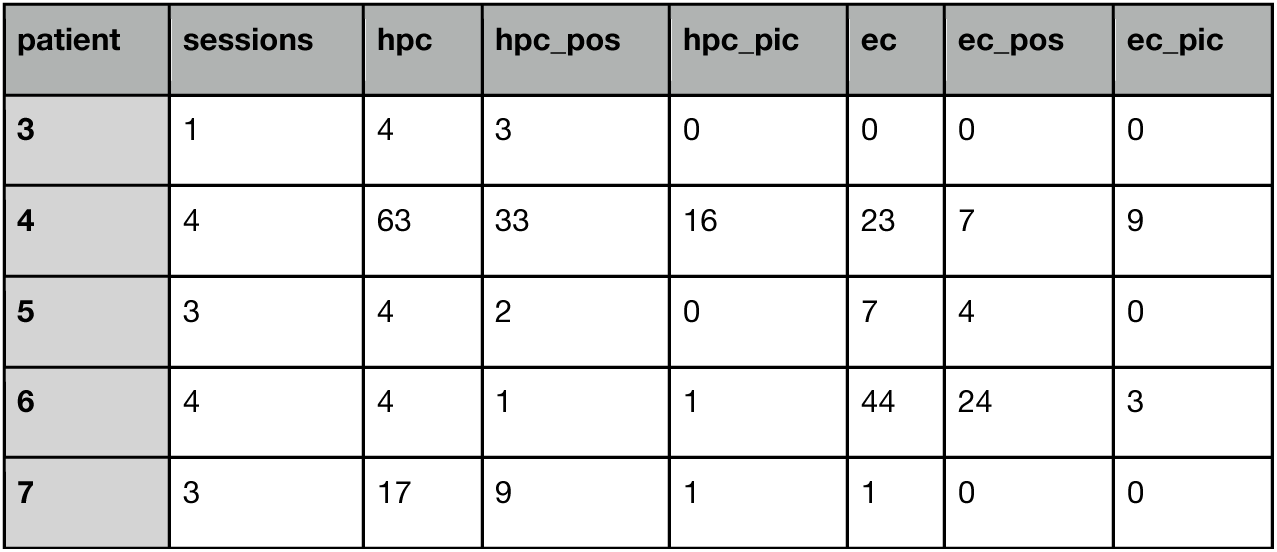
P-values table for Fig 3. P-values for orderness metric for neighbouring pairs of levels (left column), including all pairs of levels (middle column), and averaged across participants against a null distribution from shuffled level labels (right column) in dataset 1 (first row), dataset 2 (second row), and both combined (third row). The rows correspond to figure 3h-j: first row to 3h, second row to 3i, third row to 3j. The columns also correspond to figure 3h-j: first column to 3h-j left, the second column to 3h-j middle and the third column to 3h-j right. For Dataset 1, the significance threshold was corrected for multiple comparisons across both hemispheres.

**Supplementary figure 1.3.**
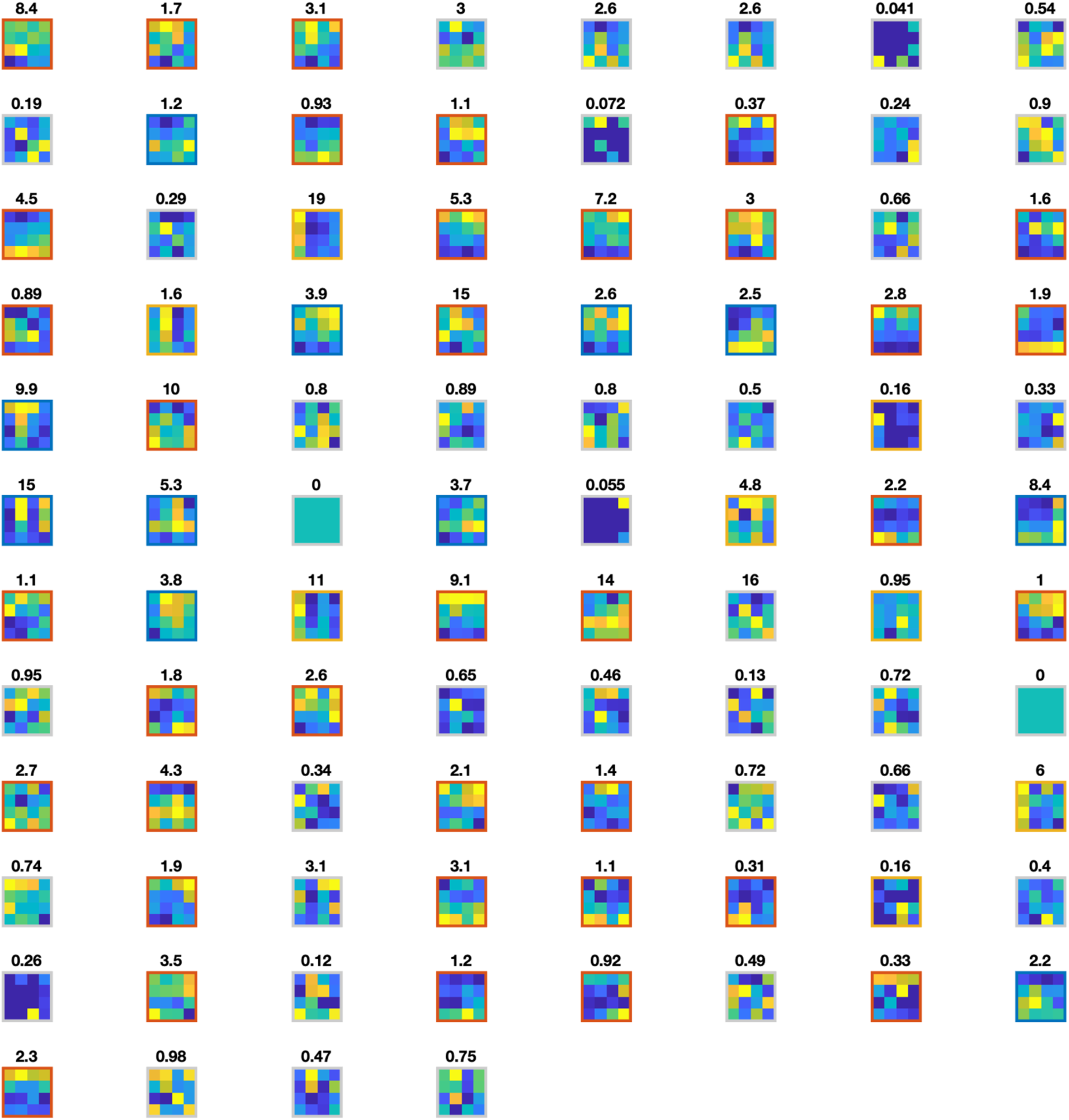
Summary firing rate plots for all hippocampal cells. For each cell we average firing rates across time and across trial stages with a particular picture (columns) in a particular position (rows), then plot the 4×4 firing rate map with the maximum firing rate in yellow and the minimum firing rate in blue. The number above each plot indicates the maximum averaged firing rate, i.e. the firing rate for the yellow square (if that number is 0, it means the cell didn’t fire during the stimulus presentation stages of the task, but it was active in different task stages). The box outline denotes whether the cell is significantly tuned to picture and position (blue), position only (orange), picture only (yellow), or neither (grey).

**Supplementary figure 1.4.**
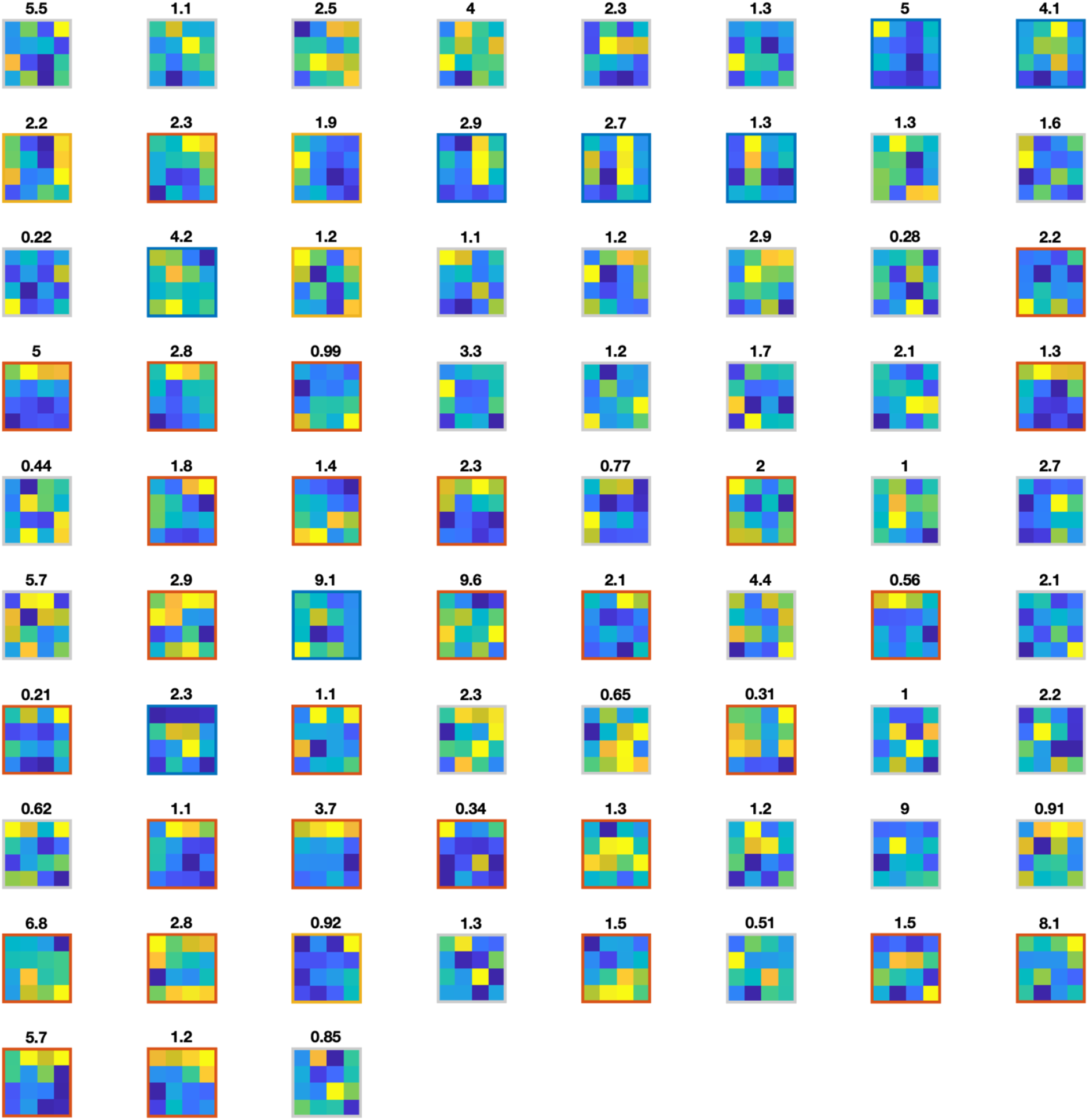
Summary firing rate plots for all entorhinal cells. For each cell we average firing rates across time and across trial stages with a particular picture (columns) in a particular position (rows), then plot the 4×4 firing rate map with the maximum firing rate in yellow and the minimum firing rate in blue. The number above each plot indicates the maximum averaged firing rate, i.e. the firing rate for the yellow square (if that number is 0, it means the cell didn’t fire during the stimulus presentation stages of the task, but it was active in different task stages). The box outline denotes whether the cell is significantly tuned to picture and position (blue), position only (orange), picture only (yellow), or neither (grey).

**Supplementary figure 1.5.**
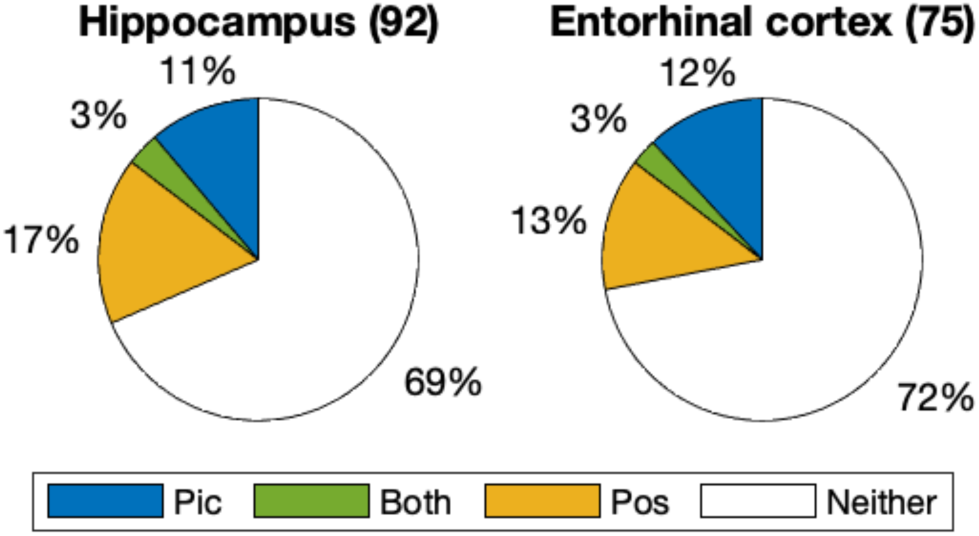
Single unit tuning to only position 2 and 3. Cells that significantly code for stimulus position, picture, or both (analogous to Fig 1m) when excluding all data from position 1 and 4.

**Supplementary figure 1.6.**
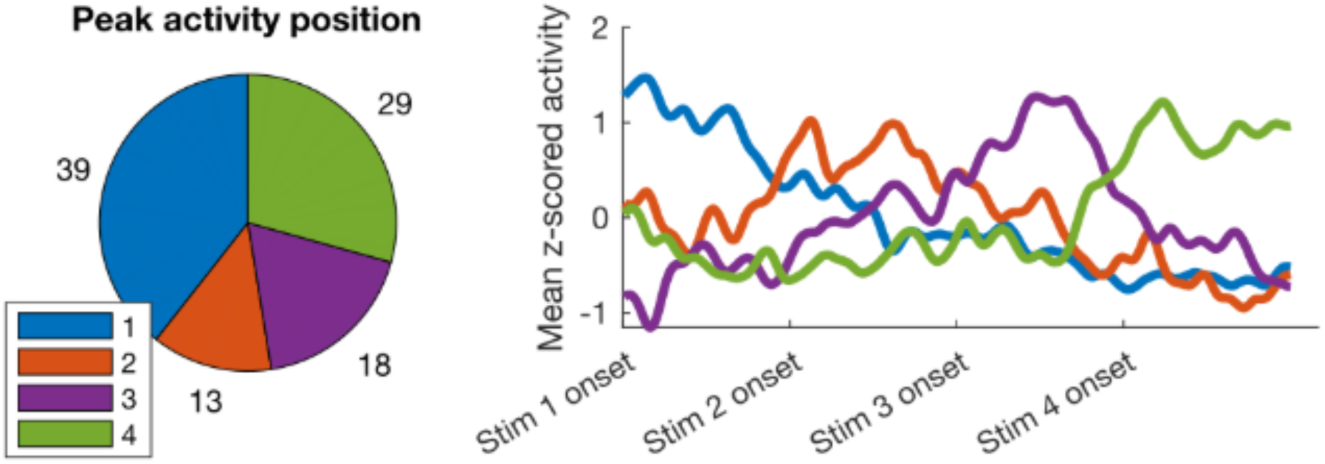
Single unit position preference. Number of cells within position coding cells that have a maximum firing rate for each position (left), with the average z-scored firing rates of each group plotted throughout the stimulus presentations stages of the trial (right).

**Supplementary figure 2.1.**
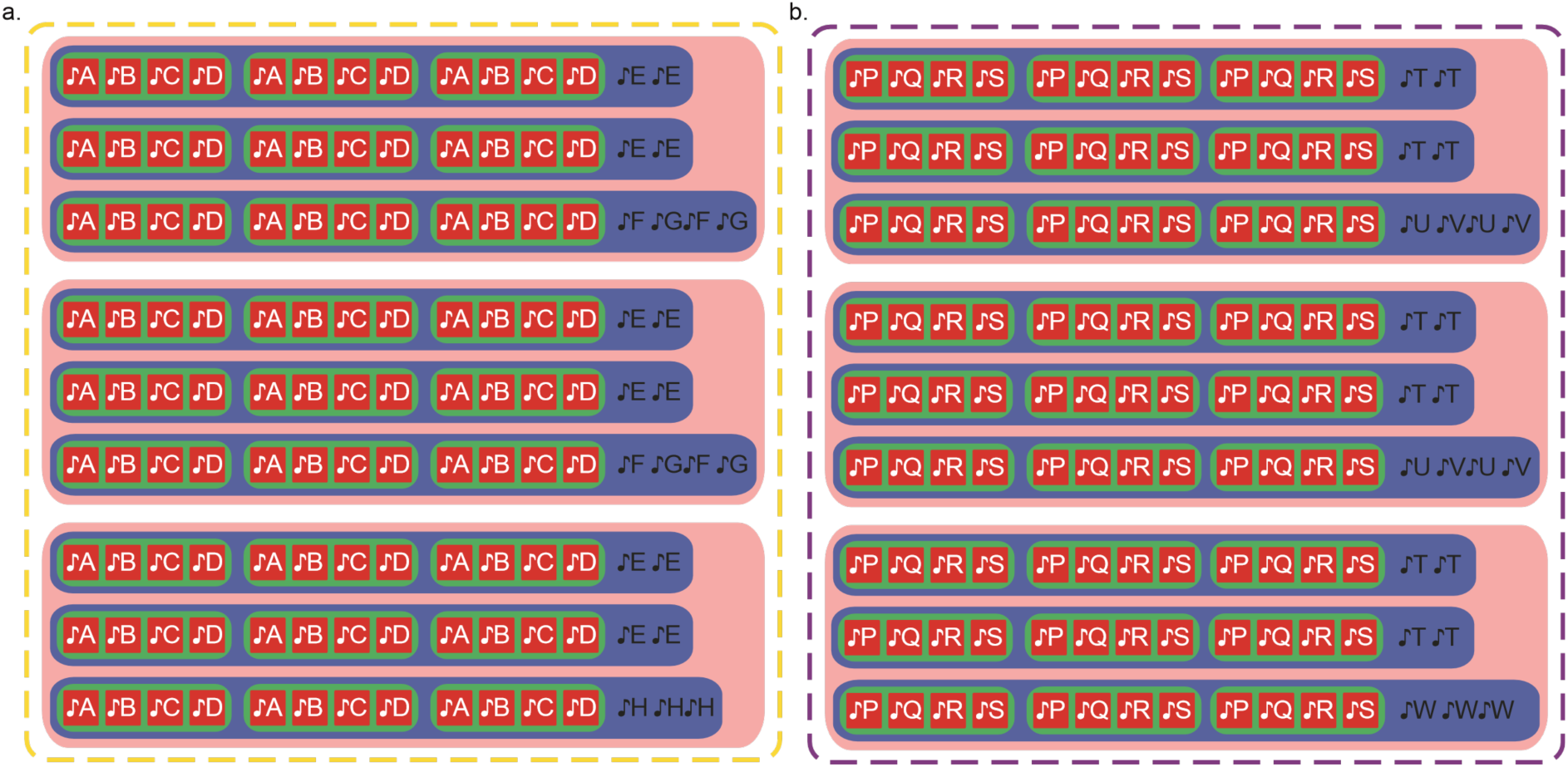
Auditory sequences. Participants learnt two sequences with identical structures but different sounds.

**Supplementary figure S2.2.**
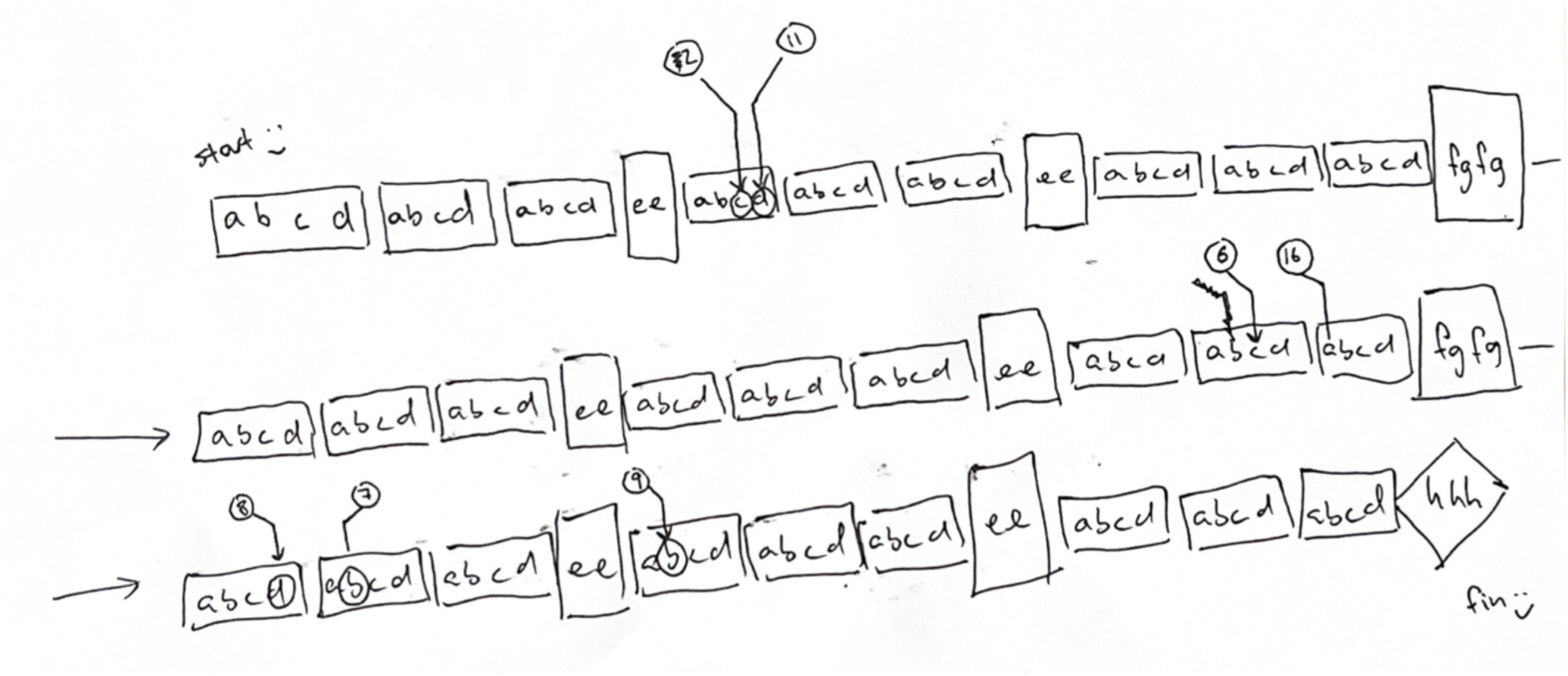
**An example participant drawing** of the auditory sequence and pictures in it. The letters are indicating the sounds in the sequence, and the numbers are referring to the pictures. We did not explicitly instruct participants about the hierarchical nature of the sequence; the division along blocks and rows is their own interpretation.

**Supplementary figure S3.1.**
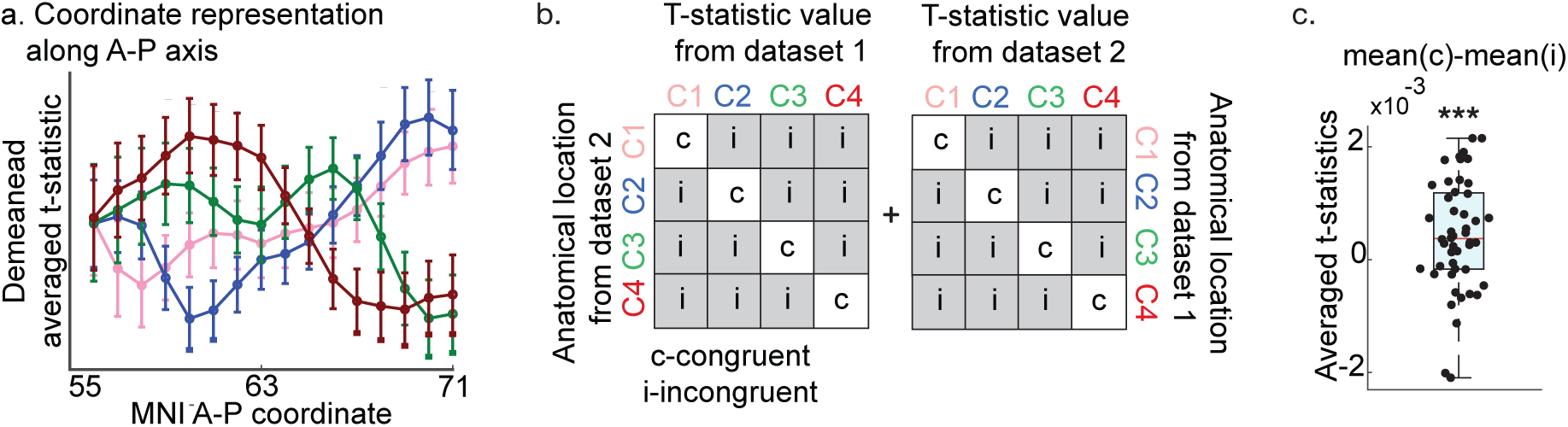
a. T-statistic for each level’s coordinate representation (C1 is pink, C2 is blue, C3 is green, C4 is red) averaged across participants within A-P slices as a function of slice A-P position in MNI space. Note that the peaks are ordered according to the hypothesised order (both datasets together). **b.** To test for the anatomical consistency of EC coordinate representations across datasets, we compare a level’s t-statistic at that level’s centre of mass from the other dataset (c: congruent) against its t-statistics at other levels centre of mass from the other dataset (i: incongruent). See Methods for more details. **c.** Congruent effects were stronger than non-congruent effects, p=0.0039.

**Supplementary Table 1.** Cells and sessions per patient. Number of sessions completed for each patient, with the result number of hippocampal (hpc) and entorhinal (ec) units and the corresponding neurons significantly tuned to position (hpc_tbls1.

| Data | Neighbours | All | Permutations |
| --- | --- | --- | --- |
| Dataset 1 | 0.00065 | 0.00002 | 0.00028 |
| Dataset 2 | 0.01911 | 0.04002 | 0.02291 |
| Both | 0.00009 | 0.00003 | 0.00004 |

**Supplementary figure S3.2.**
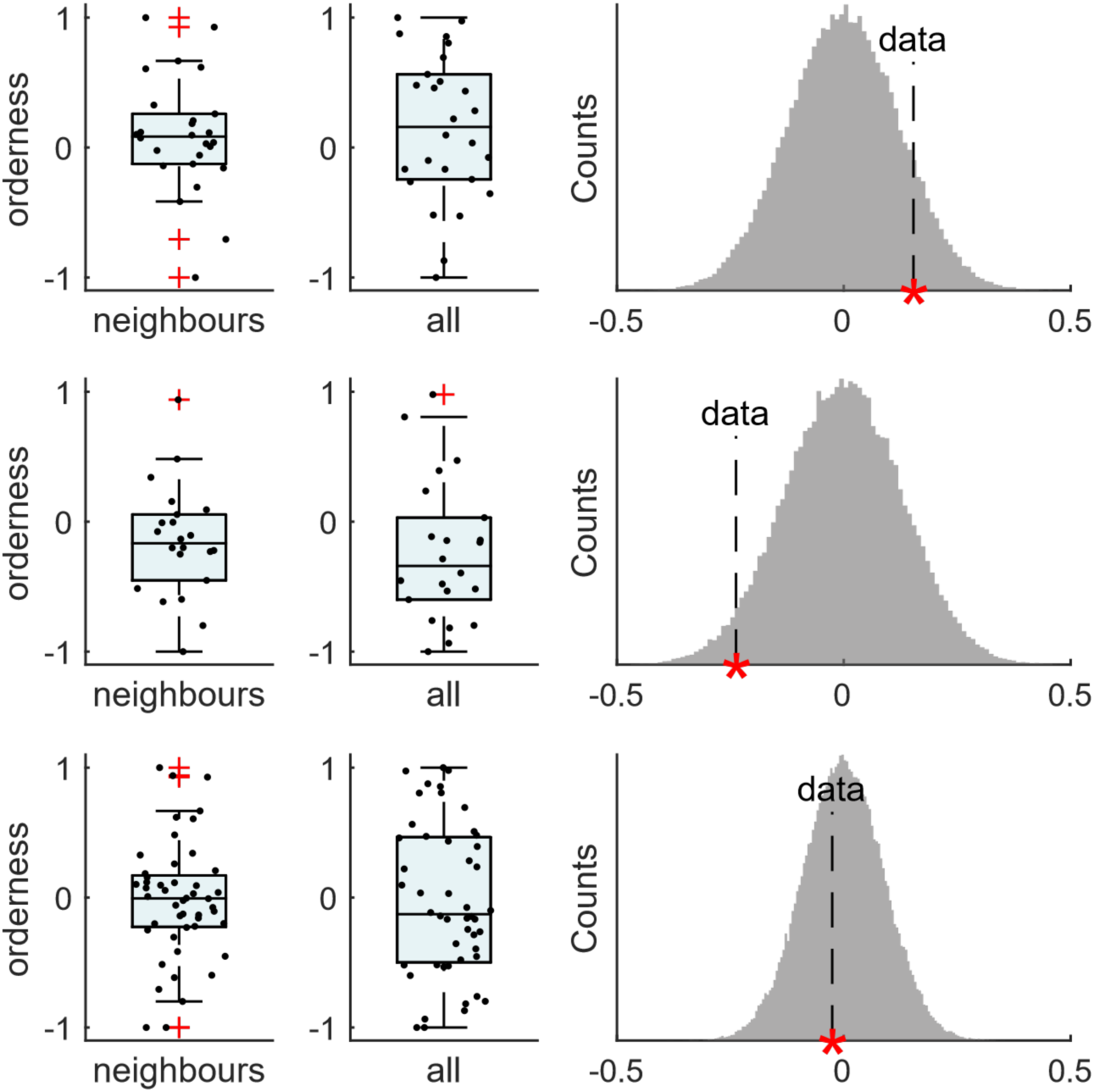
The orderness metrics computed on centres of mass defined in **the right entorhinal mask**. Orderness metric for neighbouring pairs of levels (left column), including all pairs of levels (middle column), and averaged across participants against a null distribution from shuffled level labels (right column) in dataset 1 (first row), dataset 2 (second row), and both combined (third row). See corresponding p-values in the table below.

**Supplementary Table 2.**
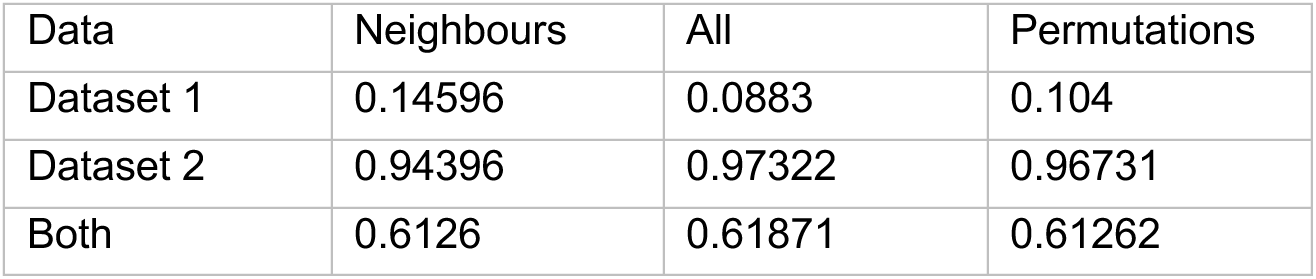
P-values table, corresponding to the Fig S3.2, for the orderness metrics computed on centres of mass defined in the right entorhinal mask. For Dataset 1, the significance threshold was corrected for multiple comparisons across both hemispheres.

| Data | Neighbours | All | Permutations |
| --- | --- | --- | --- |
| Dataset 1 | 0.14596 | 0.0883 | 0.104 |
| Dataset 2 | 0.94396 | 0.97322 | 0.96731 |
| Both | 0.6126 | 0.61871 | 0.61262 |

**Supplementary figure S3.3.**
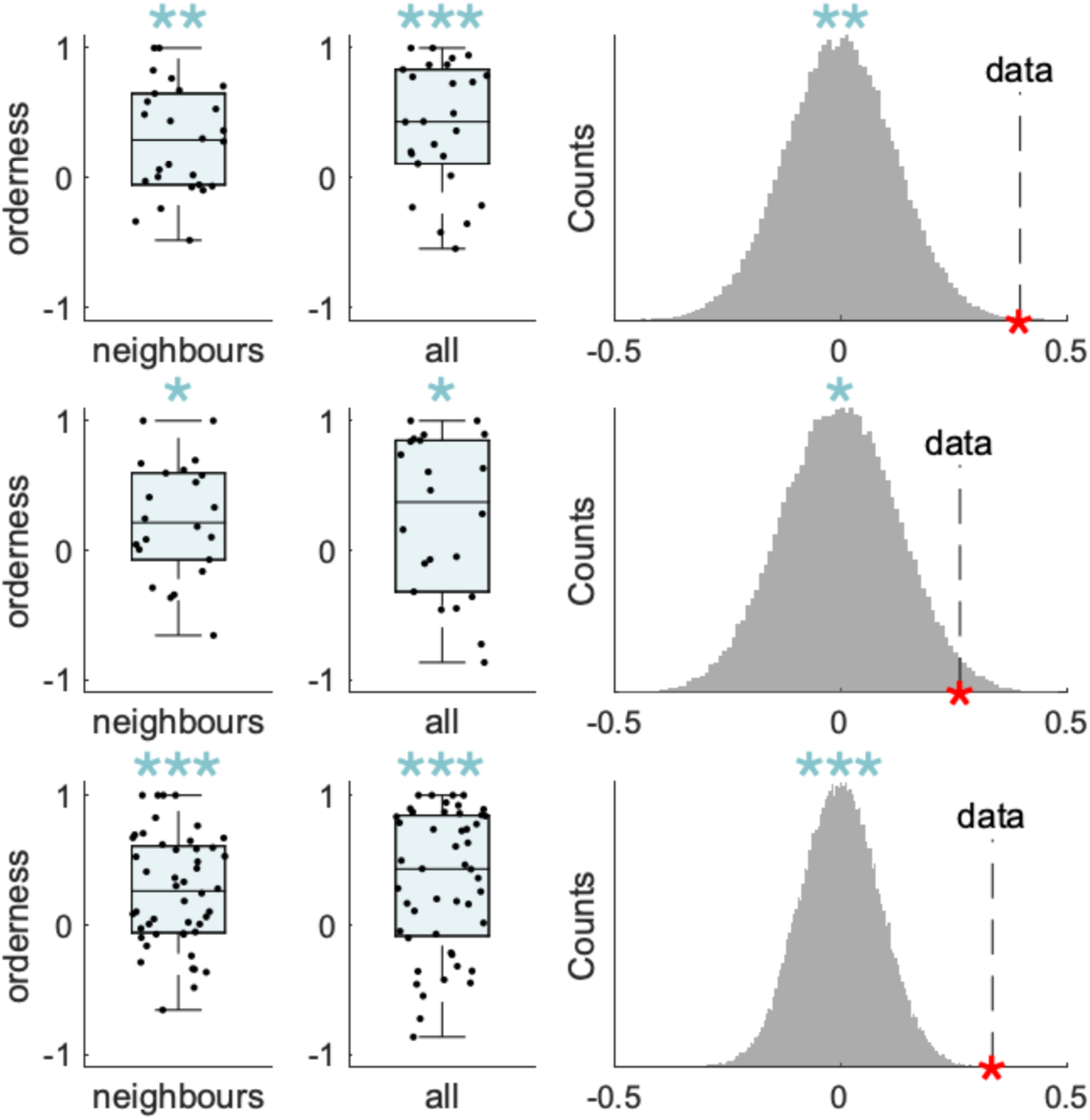
The orderness metrics computed on centres of mass defined in the **stricter entorhinal mask** (atlas probability 75% instead of the 50% probability mask used in all other analyses, see Methods). Orderness metric for neighbouring pairs of levels (left column), including all pairs of levels (middle column), and averaged across participants against a null distribution from shuffled level labels (right column) in dataset 1 (first row), dataset 2 (second row), and both combined (third row). See corresponding p-values in the table below.

**Supplementary Table 3.** P-values table, corresponding to the Fig S3.3, for the orderness metrics computed on centres of mass in the stricter entorhinal mask. For Dataset 1, the significance threshold was corrected for multiple comparisons across both hemispheres.

| Data | Neighbours | All | Permutations |
| --- | --- | --- | --- |
| Dataset 1 | 0.00093 | 0.00014 | 0.00042 |
| Dataset 2 | 0.01131 | 0.02752 | 0.02018 |
| Both | 0.00005 | 0.00004 | 0.00007 |

**Supplementary figure S3.4.**
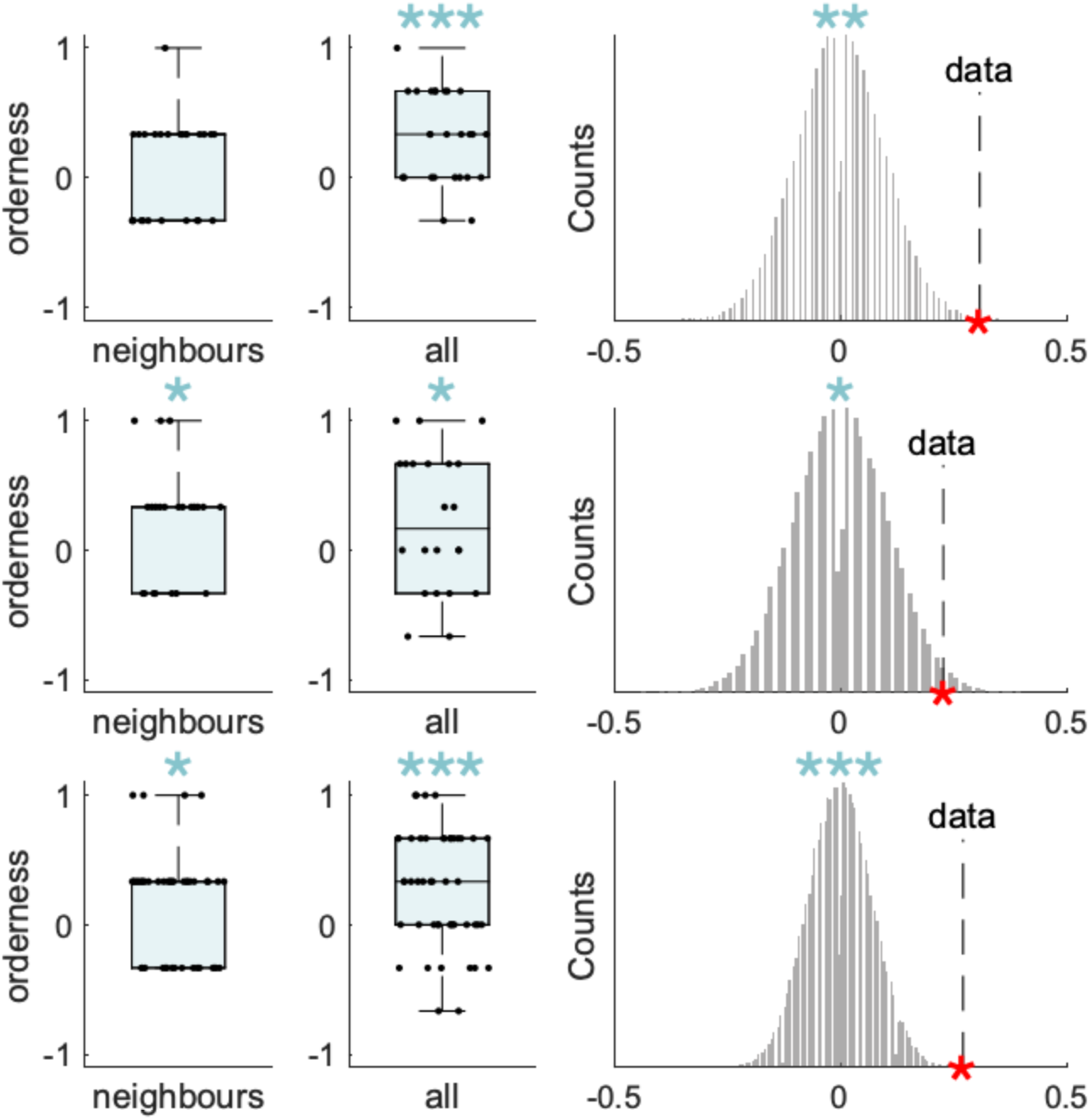
Discrete orderness metric. (see Methods) for neighbouring pairs of levels (left column), including all pairs of levels (middle column), and averaged across participants against a null distribution from shuffled level labels (right column) in dataset 1 (first row), dataset 2 (second row), and both combined (third row). See corresponding p-values in the table below.

**Supplementary Table 4.** P-values table, corresponding to the Fig S3.4, for the discrete orderness metric. For Dataset 1, the significance threshold was corrected for multiple comparisons across both hemispheres.

| Data | Neighbours | All | Permutations |
| --- | --- | --- | --- |
| Dataset 1 | 0.15639 | 0.00008 | 0.00041 |
| Dataset 2 | 0.0178 | 0.03047 | 0.01219 |
| Both | 0.01185 | 0.00005 | 0.00005 |

**Supplementary figure S3.5.**
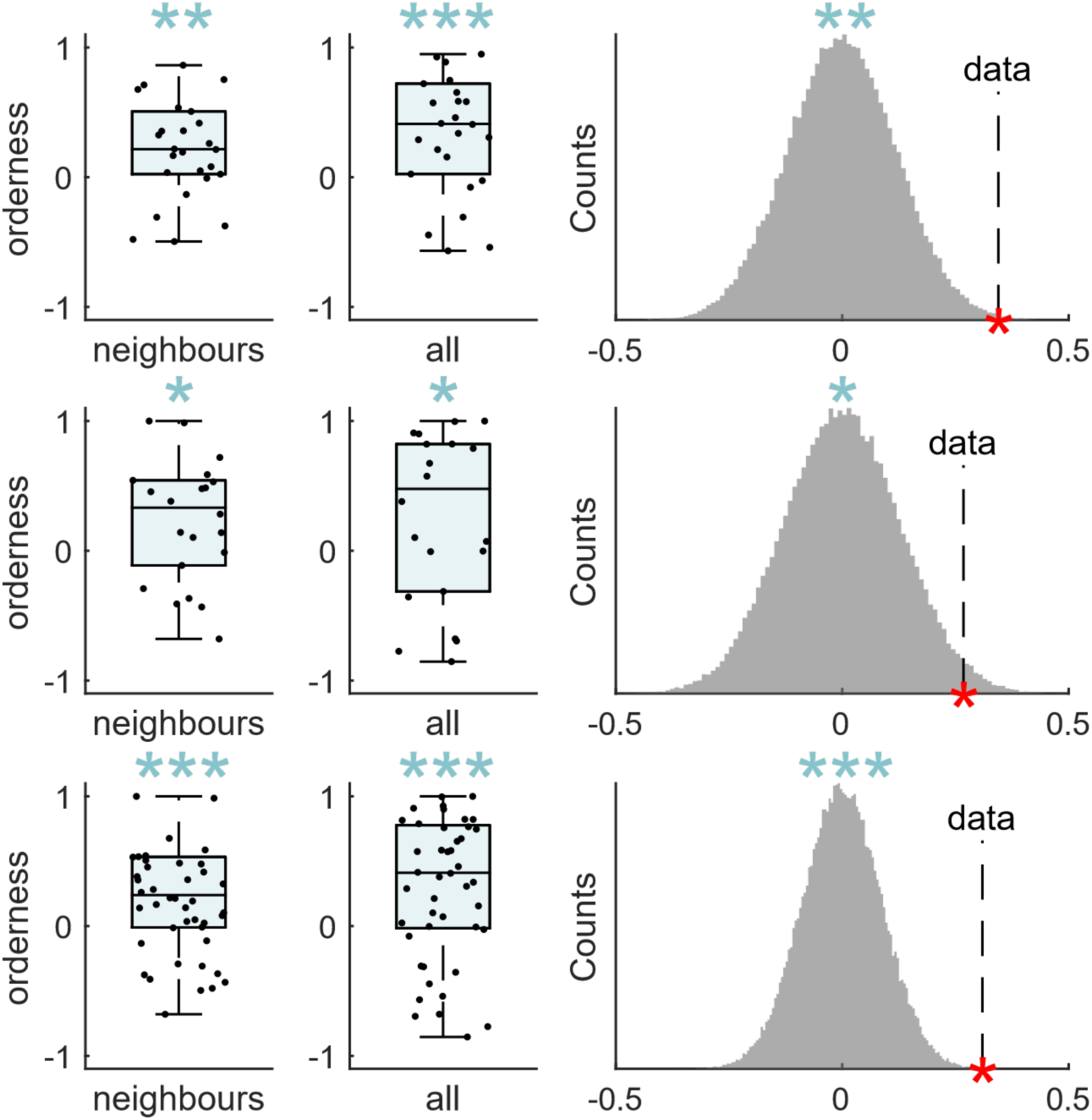
The orderness metrics computed on centres of mass defined on statistical maps that result from **separate regressions** (fitting 4 similarity regressors separately (one for each level), Methods). Orderness metric for neighbouring pairs of levels (left column), including all pairs of levels (middle column), and averaged across participants against a null distribution from shuffled level labels (right column) in dataset 1 (first row), dataset 2 (second row), and both combined (third row). See corresponding p-values in the table below.

**Supplementary Table 5.** P-values table, corresponding to the Fig S3.5, for the orderness metrics computed on statistical maps that result from a multiple regression. For Dataset 1, the significance threshold was corrected for multiple comparisons across both hemispheres.

|  | Neighbours | All | Permutations |
| --- | --- | --- | --- |
| Dataset 1 | 0.00355 | 0.00037 | 0.00173 |
| Dataset 2 | 0.01312 | 0.03182 | 0.01935 |
| Both | 0.00023 | 0.00014 | 0.00015 |

**Supplementary Table 6.** P-values table for Fig 4: across-sequence similarities matrices. P-values for orderness metric for neighbouring pairs of levels (left column), including all pairs of levels (middle column), and averaged across participants against a null distribution from shuffled level labels (right column) in dataset 1 (first row), dataset 2 (second row), and both combined (third row). The rows correspond to Fig 4f-h: first row to 3f, second row to 3g, third row to 3h. The columns also correspond to Fig 3f-h: first column to 3f-h left, the second column to 3f-h middle and the third column to 3f-h right. For Dataset 1, the significance threshold was corrected for multiple comparisons across both hemispheres.

| Data | Neighbours | All | Permutations |
| --- | --- | --- | --- |
| Dataset 1 | 0.00256 | 0.0008 | 0.00171 |
| Dataset 2 | 0.09224 | 0.05094 | 0.04066 |
| Both | 0.00171 | 0.00034 | 0.00036 |

**Supplementary figure. 3.6.**
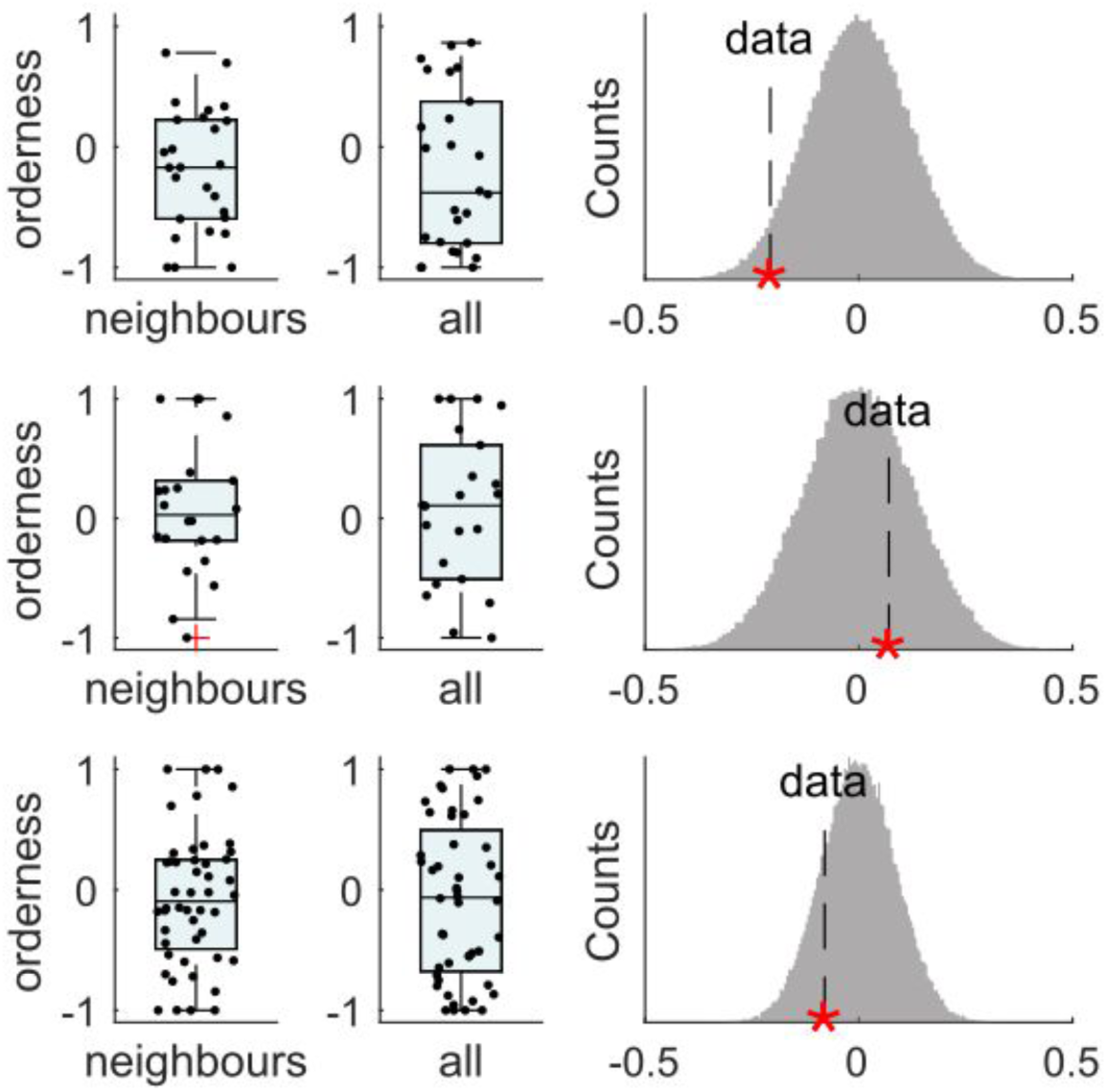
The null result for orderness metrics computed on centres of mass defined in the hippocampal mask. Orderness metric for neighbouring pairs of levels (left column), including all pairs of levels (middle column), and averaged across participants against a null distribution from shuffled level labels (right column) in dataset 1 (first row), dataset 2 (second row), and both combined (third row).

**Supplementary figure. 3.7.**
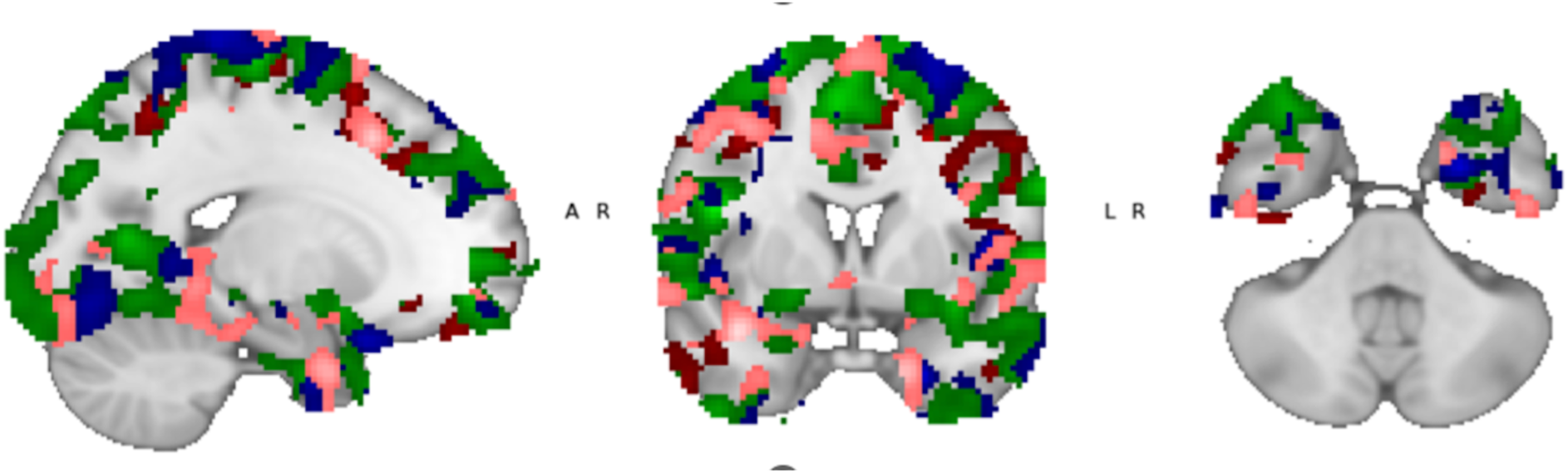
Whole-brain statistical maps for the representational similarity analysis (RSA) of each hierarchical layer coordinate. Maps are shown at a low statistical threshold for visualization purposes. Note that these group-level maps were not used for any statistical analysis. Instead, we use subject-level statistical maps to define a within-subject measure of representational order (the “orderness metric”), which is the reported statistic in all analyses. This within-subject order measure is more sensitive because it avoids signal loss due to averaging in a small brain region that is difficult to align across participants. Because widespread activation is typical in whole-brain fMRI analyses, our study used a strictly hypothesis-driven approach, focusing specifically on representational order within subjects within the entorhinal cortex.

**Supplementary figure 5.**
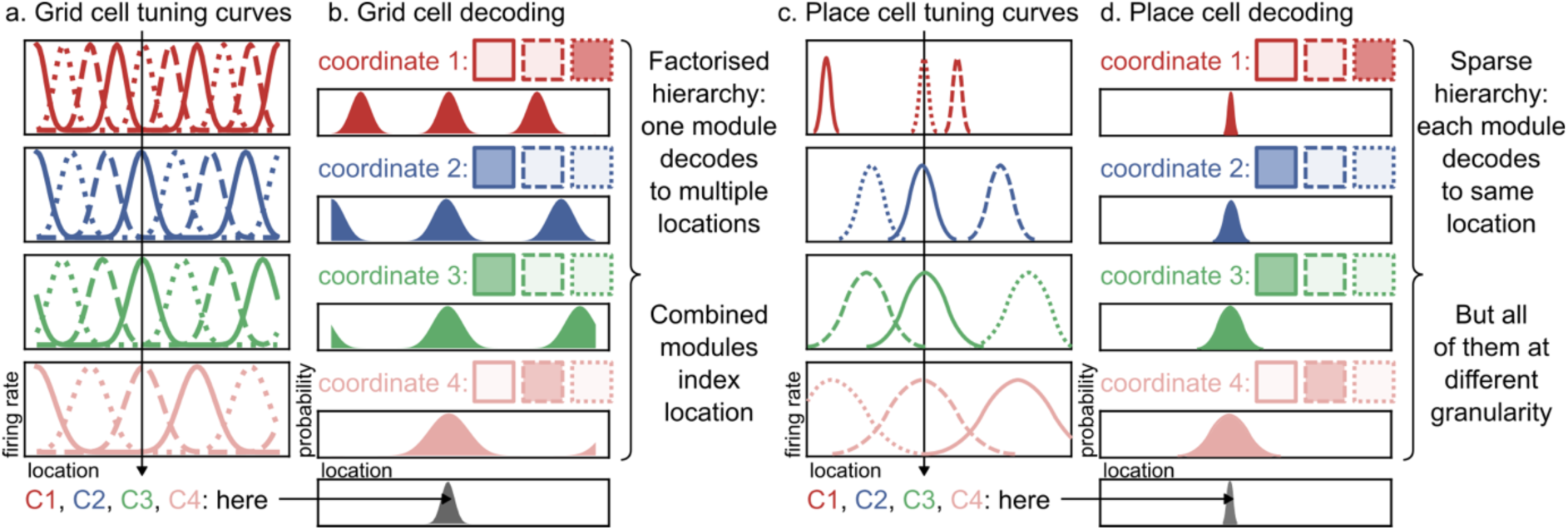
Grid cell and place cell hierarchies. a. Grid cell tuning curves, exhibiting periodic responses at different phases within a module (dotted/dashed/solid lines) at different scales across modules (colours). b. Decoded location from each module independently (colours) and jointly (grey, bottom row). These grid responses implement a factorised hierarchy, where each module encodes an independent hierarchical coordinate, and decoding a single location is only possible by combining the representations across modules – just like a calendar event is only fully determined by time, date, and month and not by any one of those. c. Place cell tuning curves that show individual peaks at different locations (dotted/dashed/solid lines) at different spatial scales (colours). d. Decoded location from each module independently (colours) and jointly (grey, bottom row). The location can be decoded separately from every module, at different levels of granularity – just like a street address that specifies street, town, and country.

